# The Spatio-Temporal Control of Zygotic Genome Activation

**DOI:** 10.1101/488056

**Authors:** George E. Gentsch, Nick D. L. Owens, James C. Smith

## Abstract

One of the earliest and most significant events in embryonic development is zygotic genome activation (ZGA). In several species, bulk transcription begins at the mid-blastula transition (MBT) when, after a certain number of cleavages, the embryo attains a particular nuclear-to-cytoplasmic (N/C) ratio, maternal repressors become sufficiently diluted, and the cell cycle slows down. Here we resolve the frog ZGA in time and space by profiling RNA polymerase II (RNAPII) engagement and its transcriptional readout. We detect a gradual increase in both the quantity and the length of RNAPII elongation before the MBT, revealing that >1,000 zygotic genes disregard the N/C timer for their activation, and that the sizes of newly transcribed genes are not necessarily constrained by cell cycle duration. We also find that Wnt, Nodal and BMP signaling together generate most of the spatio-temporal dynamics of regional ZGA, directing the formation of orthogonal body axes and proportionate germ layers.

## INTRODUCTION

The genomes of multicellular organisms are transcriptionally silent at the time of fertilisation, and the events of early development, including zygotic (also known as embryonic) genome activation (ZGA), are directed by maternal gene products (De Iaco et al., 2019; Eckersley-Maslin et al., 2019; Gentsch et al., 2018b; Lee et al., 2013; Leichsenring et al., 2013; Liang et al., 2008). The number of cell cycles after which ZGA becomes essential for development (at which embryos arrest if transcription is inhibited) is highly reproducible within each species. In the zebrafish, the frog *Xenopus* and the fruit fly *Drosophila*, this occurs after 10, 12 and 13 cell cycles, respectively, at the so-called mid-blastula transition (MBT) (Blythe and Wieschaus, 2015; Gentsch et al., 2018b; Kane and Kimmel, 1993; Newport and Kirschner, 1982a). Early development in these species occurs with no gain in cytoplasmic volume, and studies in *Xenopus* suggest that ZGA is triggered at a particular nuclear-to-cytoplasmic (N/C) ratio, when the increasing amount of nuclear DNA titrates out maternally deposited repressors (Newport and Kirschner, 1982b). Slower-developing mammalian embryos show major waves of RNA polymerase II (RNAPII)-mediated transcription as early as the 2-cell stage in mice (Bolton et al., 1984; Hamatani et al., 2004) and 4- to 8-cell stage in humans (Braude et al., 1988; Vanessa et al., 2011). This occurs days before the formation of the blastocyst, which, like the blastula, contains the pluripotent cells that form the embryo proper.

In *Xenopus*, ZGA is associated with changes in cell behaviour after the MBT. First, rapid and nearly synchronous cell cleavages give way to longer and asynchronous cell divisions (Anderson et al., 2017; Newport and Kirschner, 1982a). Second, embryonic cells acquire the ability to respond to inductive signaling (Gentsch et al., 2018b), causing them to become motile, to establish dorso-ventral patterning, and to contribute to one or two of the three germ layers (endoderm, mesoderm and ectoderm). These germ layers emerge first during gastrulation and are the primordia of all organs. Third, embryos show accelerated degradation of maternal RNA, and fourth, cells gain apoptotic (Stack and Newport, 1997) and immunogenic (Gentsch et al., 2018a) capabilities.

While large-scale ZGA occurs at the MBT, some genes escape the repressive environment of the early embryo, and nascent transcripts can be detected in *Xenopus* during rapid cleavage stages. For example, primary microRNA transcripts of the polycistronic MIR-427 gene (Lund et al., 2009) are detectable in *X. tropicalis* after just three cell divisions (Owens et al., 2016). MIR-427, like its zebrafish equivalent MIR-430, is activated at early stages by the synergistic and pioneering activities of maternal members of the SoxB1 and Pou5F (Oct4) transcription factor (TF) families (Gentsch et al., 2018b; Heyn et al., 2014; Lee et al., 2013). These core pluripotency TFs, represented by Sox3 and Pou5f3 in *Xenopus*, are characterized by ubiquitous and high translation frequencies in pre-MBT embryos (Gentsch et al., 2018b; Lee et al., 2013). Zygotic transcription of the Nodal-encoding genes *nodal3/5/6*, and of the homeobox genes *siamois1/2*, is initiated by nuclear β-catenin as early as the 32-cell stage (Owens et al., 2016; Skirkanich et al., 2011; Yang et al., 2002).

While miR-427 (and miR-430 in zebrafish) contributes to the clearance of maternal RNA (Giraldez et al., 2006; Lund et al., 2009), *nodal* and *siamois* gene initiate the formation of the germ layers and body axes (Agius et al., 2000; Lemaire et al., 1995). All these genes, and other early-activated genes in *Drosophila, Xenopus* and zebrafish, have coding sequences of <1 kb and either no introns or just a few (Heyn et al., 2014). It has been suggested that the early rapid cell cycles cause DNA replication machinery to interfere with the transcription of larger genes (Shermoen and O’Farrell, 1991), a suggestion supported, to date, by the profiling of nascent transcripts (Heyn et al., 2014). We note, however, that the detection and temporal resolution of de novo transcription can be particularly challenging for genes that have both maternal and zygotic transcripts.

Here we use the continuous occupancy of RNAPII along gene bodies as a method to record ZGA. In contrast to transcript profiling techniques, this method (1) directly determines the activity of every gene; (2) is independent of metabolic labeling (Heyn et al., 2014) and of any gene feature such as introns (Lee et al., 2013), single nucleotide polymorphisms (Harvey et al., 2013; Lott et al., 2011) or transcript half-lives; and (3) circumvents difficulties in detecting nascent transcripts in a large pool of maternal transcripts. By combining these data with the profiling of the transcriptome along the primary body axes (Blitz et al., 2017), we resolve ZGA in time and space for wild-type and various loss-of-function embryos. We provide evidence that runs counter to our original understanding of the cell cycle or of the N/C ratio in constraining gene expression before MBT. And finally, we show how signaling initiates and coordinates spatio-temporal ZGA in the *Xenopus* embryo.

## RESULTS

### RNAPII Profiling Reveals Exponential ZGA before MBT

In an effort to resolve the progression of ZGA, we profiled chromatin for RNAPII engagement on hand-sorted *X. tropicalis* embryos over six developmental stages from the 32-cell to the late gastrula stage (Figure 1A,B). RNAPII was localized on the genome by chromatin immunoprecipitation followed by deep sequencing (ChIP-Seq). We complemented RNAPII profiling with high time-resolution transcriptomics (Owens et al., 2016) counting both exonic and intronic RNA at 30-min intervals from fertilization to the late gastrula stage (Figures 1A,B and S1A,B). For both maternal and zygotic genes, the detection threshold was set to ≥3 transcripts per million (TPM) averaged over any 1-h window during this developmental period to avoid genes with general low-level expression. This restricted the analysis to 13,042 genes (Figure 1B). These genes were considered active when we detected simultaneously RNAPII enrichment along their full length (see Transparent Methods) as well as the presence of the corresponding transcripts. In doing so, we used a low threshold of ≥0.1 TPM so as not to miss the onset of gene transcription. RNAPII-guided ZGA profiling was verified in part by active post-translational histone marks (Hontelez et al., 2015) and by differential expression methods aiming at detecting nascent transcripts. Thus, zygotic transcript depletion (by blocking RNAPII elongation with α-amanitin) (Gentsch et al., 2018b) or enrichment (by selecting 4-thiouridine [4sU] tagged transcripts at the MBT and the mid-gastrula stage) showed substantial overlaps and positive correlations with RNAPII-covered genes (Figures 1A,B and S1C,D and Tables S1 and S2).

**Figure 1.**
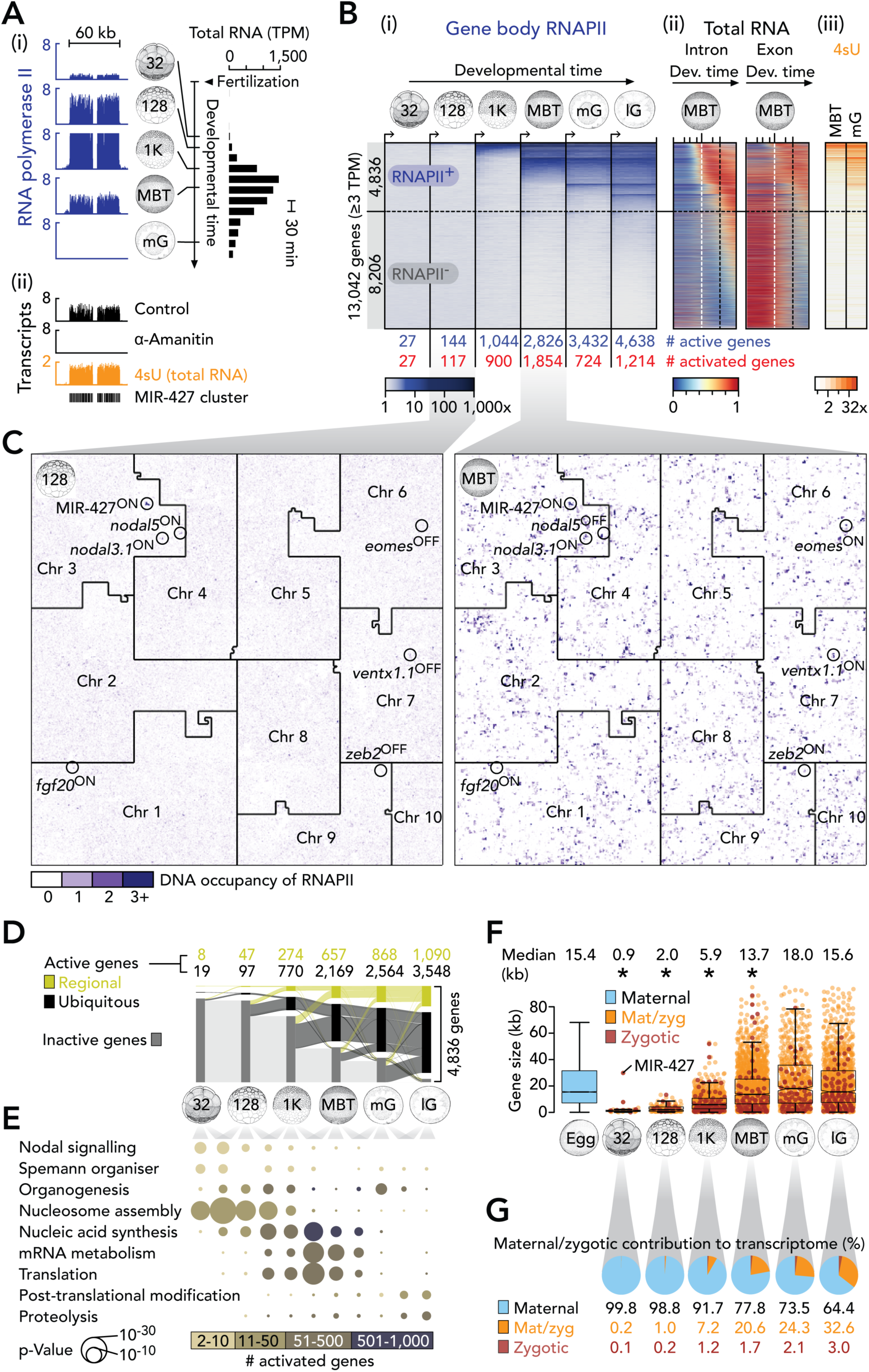
Dynamics and Architecture of ZGA in *X. tropicalis*. (A) (i) Genome-wide profiling of RNAPII and total RNA (Owens et al., 2016) to determine temporal ZGA dynamics. (ii) Complementary approach to detect transcriptionally active genes by α-amanitin-induced loss and 4sU-enrichment of nascent (zygotic) transcripts. (B) Progression of ZGA from the 32-cell to the late gastrula stage based on (i) whole gene body (full-length) occupancy of RNAPII (i.e., RNAPII was enriched across entire gene bodies; see Transparent Methods). Co-aligned: (ii) High time-resolution of total RNA, separated by intron- and exon-derived signals, from fertilization to the late gastrula stage, and (iii) enrichments of 4sU-tagged RNA at the MBT and the mid-gastrula stage. Numbers below RNAPII heat map represent counts of active (blue) and activated (red) genes at the indicated developmental stages. The horizontal dotted line separates RNAPII-engaged (RNAPII^+^) from non-engaged (RNAPII^−^) genes as detected until the late gastrula stage. The vertical dotted lines in the total RNA plots indicate the developmental time points of the MBT (white) and the late gastrula stage (black), respectively. (C) 2D space-filling (Hilbert) curves showing RNAPII recruitment to chromosomes (Chr) at the 128-cell stage and the MBT. A few zygotic genes are highlighted as being active (ON) or not (OFF) based on their engagement with RNAPII. (D) Alluvial diagram of spatio-temporal ZGA. Tissue-specificity inferred from regional transcript enrichments along the animal-vegetal or the dorso-ventral axes or both (Blitz et al., 2017). (E) ZGA-associated enrichment of biological processes. (F) Length of maternal and/or zygotic genes. *, p <1.9e-7 (Wilcoxon rank-sum test against maternal and post-MBT activated genes); r_effect_ 0.06 (‘MBT’ vs ‘Egg’) - 0.48 (‘128’ vs ‘mG’). (G) Maternal/zygotic contribution to the transcriptome deduced from full-length RNAPII occupancy and maternally inherited RNA. Abbreviations 32, 32-cell; 128, 128-cell; 1K, 1,024-cell; MBT, mid-blastula transition; mG, mid-gastrula; lG, late gastrula; 4sU, 4-thiouridine; Mdn, median; TPM, transcripts per million. See also Figure S1 and Tables S1 and S2.

This analysis revealed an exponential ZGA before the MBT with 27, 144 and 1,044 active genes after 5 (32-cell, ∼2.5 hpf), 7 (128-cell, ∼3 hpf) and 10 (1,024-cell, ∼4 hpf) cell cycles, respectively. Gene activation reached its peak at the MBT (∼4.5 hpf), with 1,854 newly-activated genes, before dropping to 724 genes at the early-to-mid gastrula stage (∼7.5 hpf) and increasing again to 1,214 genes towards the end of gastrulation (∼10 hpf) (Figures 1B,C and S1E and Table S2). The dramatic increase in transcriptional activity that occurs in the 1.5 hours between the 128-cell stage and the MBT can be illustrated by Hilbert curves (Figure 1C), which provide a genome-wide overview of RNAPII enrichment by folding chromosomes into two-dimensional space-filling and position-preserving plots (Gu et al., 2016). While most zygotic genes remain active beyond the mid-gastrula stage, 197 (including *siamois2* [*sia2*], *nodal5* and *znf470*) of the 4,836 zygotic genes (∼4%) are deactivated within ∼6 h of development (Figures 1C and S1F,G). Slightly less than one third of the activated genes were differentially expressed along either or both of the animal-vegetal and dorso-ventral axes (Figures 1D and S1F).

The temporal order of enriched biological processes supported by ZGA matched the regulatory flow of gene expression, starting with nucleosome assembly, nucleic acid synthesis, mRNA metabolism and production, post-translational modification and degradation of proteins (Figure 1E). The earliest transcriptional engagement, beginning at the 32 to 128-cell stages, was detected in gene clusters of tens to hundreds of kilobases (Figures 1A,C and S1H-J). These clusters featured close relatives of the same genes, some of which are critical to Nodal signaling (Nodal ligands), the formation of the Spemann organiser (Siamois homeobox transcription factors), nucleosome assembly (histones), mRNA decay (MIR-427), and ongoing gene regulation (zinc finger [ZF] transcription factors with on average 10 Cys2-His2 [C2H2] domains; Figure S1J). These earliest activated genes were shorter and encoded smaller proteins than those within the maternal pool or those that are activated post-MBT (Figures 1F and S1K). The non-coding features that contributed most to the differences in length were the 3’ UTRs and introns (Figure S1K).

We noted that the shorter zygotic genes observed before the MBT did not correlate strictly with the time constraints imposed by short cell cycles. We detected increasing and wider spread of de novo recruitment of RNAPII before the MBT, when cleavages continue to occur at rapid and near-constant pace (Figures 1F and S1K). During this period, the median length of activated genes (and their coding sequences) increases from ∼0.9 kb (∼0.4 kb) to ∼5.9 kb (∼0.9 kb). However, it was not until after the MBT that the overall architecture of zygotic and maternal genes became indistinguishable (Figures 1F and S1K). Temporal comparison of RNAPII engagement and total RNA profiling suggested that the zygotic contribution to the transcriptome (as calculated by the number of zygotic genes divided by the number of genes with ≥0.1 TPM maternal transcripts averaged over the first hour post-fertilisation when the entire zygotic genome is still transcriptionally inactive) rose within seven cell cycles from ∼0.2% at the 32-cell stage to ∼22% at the MBT (Figure 1G). Further maternal degradation and more moderate transcriptional engagement extended the zygotic contribution to about one third of the transcriptome by the late gastrula stage. Maternal transcripts (≥0.1 TPM, see above) were detected for ∼67% of newly activated genes (18 out of 27 genes) at the 32-cell stage, ∼85% (99/117) at the 128-cell stage, ∼87% (780/900) at the 1,024-cell stage, ∼95% (1,754/1,854) at the MBT, ∼89% (644/724) at the mid-gastrula stage and ∼90% (1,094/1,214) at the late gastrula stage (Table S2). Altogether ∼91% (4,389/4,836) of newly activated genes have ≥0.1 TPM maternal contribution.

### Wnt, Nodal and BMP Signals Are Key Drivers of Regional ZGA

We next sought to investigate the single and combined effects of different inductive signals on the spatio-temporal dynamics of ZGA. The early vertebrate embryo employs canonical Wnt, Nodal and BMP signals and their key transcriptional effectors β-catenin, Smad2 and Smad1, respectively, to establish the primary body axes and the three germ layers (reviewed by Arnold and Robertson, 2009 and Kimelman, 2006). In *Xenopus*, β-catenin first translocates to the nuclei of dorsal blastomeres at the 32-cell stage (Larabell et al., 1997; Schneider et al., 1996) (Figure 2A). After the MBT, zygotic Wnt8a causes more nuclear β-catenin to accumulate around the forming blastopore lip (Christian and Moon, 1993; Schohl and Fagotto, 2002). The nuclear translocation of Smad1 and Smad2 is triggered around the MBT by various BMP and Nodal ligands. Nuclear Smad1 is primarily detected on the ventral side and the blastocoel roof of the embryo while nuclear Smad2 is detected within the vegetal hemisphere (VH) and the marginal zone (MZ) (Faure et al., 2000; Schohl and Fagotto, 2002) (Figure 2A).

**Figure 2.**
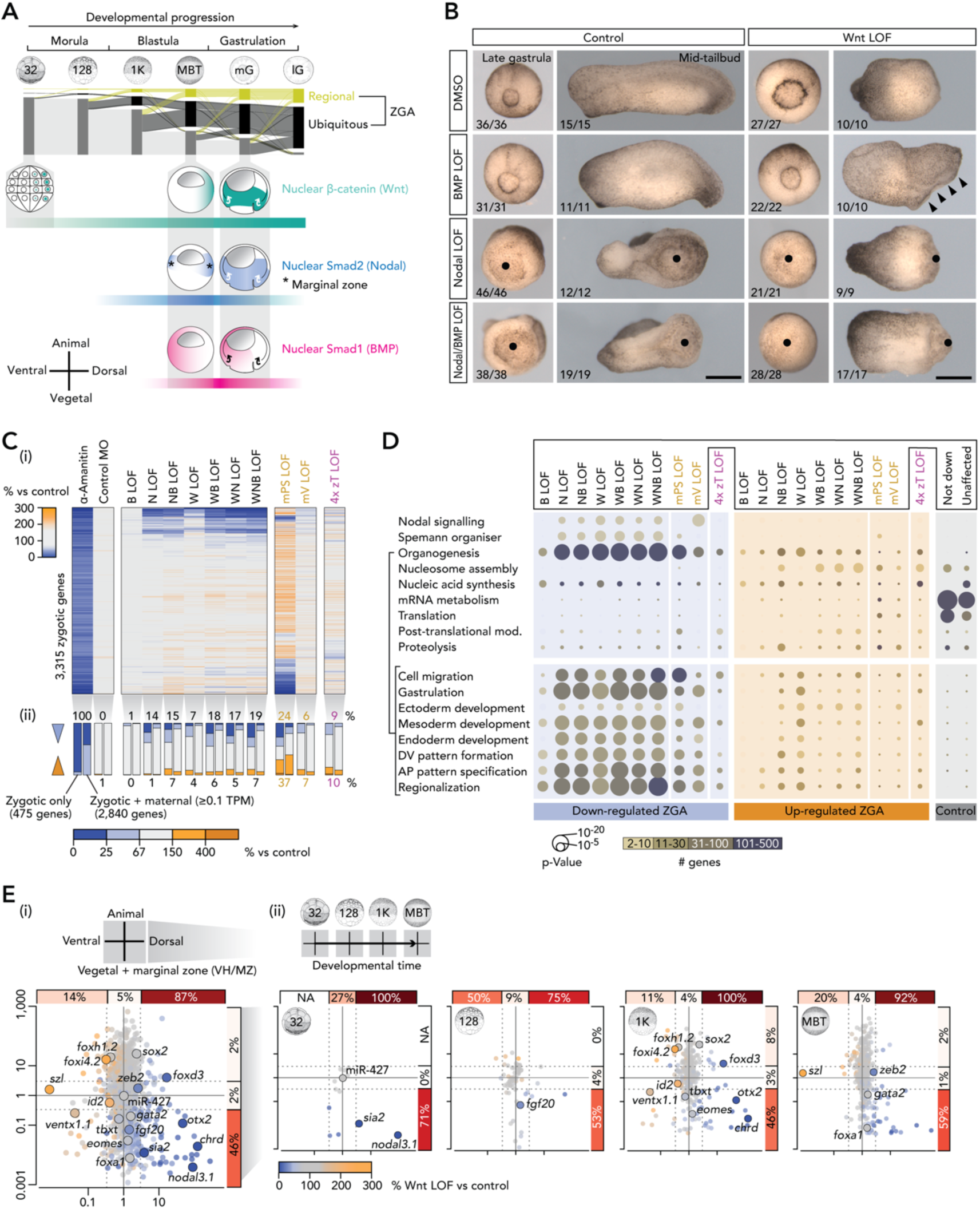
Spatio-temporal ZGA Regulated by Canonical Wnt, Nodal and BMP Signals. (A) Spatio-temporal ZGA and nuclear localization of signal mediators β-catenin (canonical Wnt), Smad2 (Nodal) and Smad1 (BMP) (Faure et al., 2000; Larabell et al., 1997; Schohl and Fagotto, 2002) from the 32-cell to the late gastrula stage. (B) Morphological phenotypes of single and combined signal LOFs at the late gastrula and the mid-tailbud stage. Left (‘control’) pictures are taken from Gentsch et al. (2018b). Bullet points, failed blastopore formation. Arrowheads, excessive neural fold formation. Scale bar, 0.5 mm. (C) Heat map (i) and bar graph summary (ii) of ZGA mis-regulated in various LOF embryos: α-amanitin, positive control; control MO, negative control. Abbreviations: B, BMP; N, Nodal; W, canonical Wnt; mPS, maternal Pou5f3/Sox3; mV, maternal VegT; 4x zT, four zygotic T-box TFs (zygotic VegT, Eomes, Tbxt and Tbxt.2). (D) Biological processes enriched with mis-regulated and control (not down-regulated or unaffected by the loss of maternal TFs or signaling) sets of zygotic genes under indicated LOFs. (E) Summary (i) and temporal resolution (ii) of Wnt LOF effects on regional ZGA. Percentages only refer to the down-regulated genes among all zygotic genes with similar expression ratios along the animal-vegetal or the dorso-ventral axes. See also Figure S2, Tables S1 and S3, and Movie S1.

In an effort to inhibit canonical Wnt signaling, we injected into the *X. tropicalis* zygote a previously validated antisense morpholino oligonucleotide (MO) which interferes with β-catenin protein synthesis by annealing to the translation start codon (Heasman et al., 2000). Nodal and BMP signals were selectively blocked by incubating dejellied embryos from the 8-cell stage in the cell-permeable inhibitors SB431542 (Ho et al., 2006; Inman et al., 2002) and LDN193189 (Cuny et al., 2008; Young et al., 2017), respectively. The morphological phenotypes of these single loss-of-function (LOF) treatments were consistent with previous observations and ranged from impaired axial elongation causing the loss of tail structures (BMP, Reversade et al., 2005) to severe gastrulation defects (Wnt, Heasman et al., 2000, and Nodal, Ho et al., 2006) as shown in Figure 2B. Briefly, Nodal LOF impaired blastopore lip formation and bulk tissue movements of gastrulation (bullet points in Figure 2B). However, it did not preclude subsequent elongation of the antero-posterior axis. By contrast, Wnt LOF embryos underwent gastrulation (albeit delayed and more circumferentially rather than in a dorsal to ventral wave), but failed to form an antero-posterior axis, with both head and tail being absent. With respect to the joint effects of Wnt, Nodal and BMP signaling, most dual and triple LOFs combined their individual morphological defects such that, for example, Wnt/Nodal LOF resulted in the complete loss of gastrulation and axial elongation. By contrast, Wnt/BMP LOF produced defects such as non-fusing neural folds (arrowheads in Figure 2B), structures that were either absent in Wnt LOF embryos or normal in BMP LOF embryos.

Changes to ZGA caused by the single or combined LOF of Wnt, Nodal and/or BMP were then determined at the late blastula stage on a transcriptome-wide scale using deep RNA sequencing (RNA-Seq). Analysis was limited to the 3,315 zygotic genes for which spatio-temporal expression data is available (Blitz et al., 2017; Owens et al., 2016) and where reduced expression (≥50% loss of exonic and/or intronic transcript counts, FDR ≤10%) could be detected in α-amanitin-injected embryos (Figure 2C and Table S3) (Gentsch et al., 2018b). α-Amanitin-mediated inhibition of RNAPII elongation impedes the morphogenetic tissue movements of gastrulation and ultimately leads to early embryonic death (Gentsch et al., 2018b). Spatial gene expression patterns were inferred from experiments comparing the transcriptomes of embryos dissected along their animal-vegetal and dorso-ventral axes (Blitz et al., 2017); we did not include the left-right axis because there were no significant differences in gene expression across this axis at the gastrula stage (Blitz et al., 2017). The signal-mediated transcriptional effects (1.5-fold change from control RNA level) on zygotic genes, 86% (2840/3315) of which have ≥0.1 TPM maternal contribution, ranged from ∼1.5% (∼1.3% down and ∼0.2% up) to ∼26% (∼19% down and 7% up) for single BMP LOF and triple Wnt/Nodal/BMP LOFs, respectively (Figure 2C). As expected, the transcript levels of genes whose expression is solely zygotic were more strongly affected than those of zygotic genes with maternally contributed transcripts (Figures 2C and 4A). The extent of ZGA mis-regulation largely reflected the severity of the resulting morphological phenotypes at the late gastrula and the mid-tailbud stage (Figure 2B,C).

In comparison, the LOFs of critical maternal TFs like Pou5f3/Sox3 or VegT (Gentsch et al., 2018b) caused the mis-regulation of 61% (∼24% down and ∼37% up) and 13% (∼6% down and ∼7% up) of zygotic genes, respectively. The LOFs of four zygotic T-box TFs (zVegT, Eomes, Tbxt and Tbxt.2), all of which require Nodal signaling for their expression, caused slight mis-regulation in 19% (∼9% down and ∼10% up) of the zygotic genes as detected over three consecutive developmental time points during gastrulation (Table S3). Among the ZGA-enriched biological functions (Figure 1D), Wnt, Nodal and BMP signals, like maternal Pou5f3/Sox3 and VegT, strongly affected zygotic genes associated with cell migration, gastrulation, dorso-ventral and antero-posterior body axis formation and regionalization (Figure 2D). Impaired tissue movements during gastrulation, as observed in various LOFs (Figure 2B and Movie S1), was prefigured by a strong enrichment for cell migration-associated genes. The genes suppressed or unaffected by the selected signals and maternal TFs were enriched for the ZGA-critical biological processes of mRNA metabolism and translation. For instance, the transient activation of the entire zinc finger cluster (Figure S2A) was not affected by any tested LOF. Because family members are frequently cross-regulated, and the MBT-staged chromatin contains many Krüppel-like zinc finger ‘footprints’ (Gentsch et al., 2018b), it is conceivable that the unaffected, tissue-nonspecific part of ZGA is regulated by maternal zinc finger TFs. This vertebrate gene regulatory branch may be more ancient than that of Pou5F/SoxB1 as zinc finger TFs like Zelda are also key to ZGA of the invertebrate *Drosophila* (Liang et al., 2008).

Next, signal-dependent ZGA was resolved in time and space based on (i) the profiling of RNAPII-engaged chromatin from the 32-cell stage to the MBT and (ii) known gene expression patterns along the animal-vegetal and dorso-ventral axes (Blitz et al., 2017) (Figures 2E and S2B-F). In line with the nuclear translocation of their signal mediators (Figure 2A), Wnt, Nodal and BMP proved to be required for gene activation in different spatio-temporal domains of the early embryo: β-catenin was needed for ∼87% and ∼46% of genes preferentially expressed on the dorsal side and in the VH/MZ, respectively. Some of its target genes like *nodal3.1* and *sia2* were already active by the 32-cell stage (Figures 1C and 2E). Upon Wnt LOF, the early transcriptional down-regulation was followed by the mis-regulation of opposing cell fate specifiers: the upregulation of ventral genes (e.g., *id2, szl*) and the downregulation of dorsal genes (e.g., *chrd, otx2*). These observations suggest that β-catenin protects dorsal cells from ventralization (Figures 2E and S2C).

Along similar lines, Nodal LOF embryos predominantly displayed a down-regulation of dorsal (∼63%) and VH/MZ-specific (∼73%) genes, although there was no effect of Nodal LOF on the earliest-activated genes at the 32-cell stage or on opposing cell fate regulators (Figure S2A,B). Among the first genes to be sensitive to Nodal LOF was the MZ-specific FGF ligand *fgf20*, activated by the 128-cell stage (Figures 1C and S2A,B). By contrast, BMP LOF caused a decrease in ventrally-expressed gene expression (∼45%) from the 1,024-cell stage onwards (Figure S2A,C).

As a comparison, the ubiquitous expression of maternal Pou5f3/Sox3 was required for transcription in all spatio-temporal domains, including, for example, the uniform expression of miR-427 (Figure S2A,D). The requirement for Pou5f3/Sox3 was more marked, however, for region-specific genes, in particular those expressed within animal-(∼55%) and ventral-specific (∼67%) domains (Figure S2A,D). The maternal TF VegT promoted vegetal identity by activating ∼40% of genes transcriptionally enriched within its own expression domain, the vegetal hemisphere, while suppressing genes that are expressed in the animal hemisphere like *foxi4.2*. The requirement for VegT was similar in ventral- and dorsal-specific genes (∼31% and ∼30% respectively).

### Wnt/BMP Synergy Enables Uniform ZGA Across the Dorso-Ventral Axis

The relationships between inductive signals with respect to spatial aspects of ZGA was explored by comparing zygotic transcriptomes in single and double LOF experiments. Interestingly, while simultaneous loss of both Nodal and BMP function, or both Nodal and Wnt, caused additive effects on gene expression compared to the single LOFs (Figures 3E and S3A-N), simultaneous loss of both BMP and Wnt signaling showed a more synergistic effect (Figure 3A,F). These observations are consistent with the morphological consequences of single and double LOFs (Figure 2B). Single LOF experiments revealed very little overlap between Wnt and BMP gene targets (Figure 3B), a result consistent with their domains of activity which are, initially, at opposite ends of the dorso-ventral axis. However, dual Wnt/BMP inhibition increased the number of downregulated genes by 292, a rise of ∼118% and ∼664% with respect to individual Wnt- and BMP-dependent genes respectively (Figure 3B). Interestingly, this synergy affected 166 Nodal-dependent genes, most of which had uniform expression levels across the dorso-ventral axis (Figure 3C,D,F-H). Thus, spatially-restricted Wnt, BMP and Nodal signals act together to establish dorso-ventral expression uniformity of genes such as *tbxt* and *eomes* (Figure 3I).

**Figure 3.**
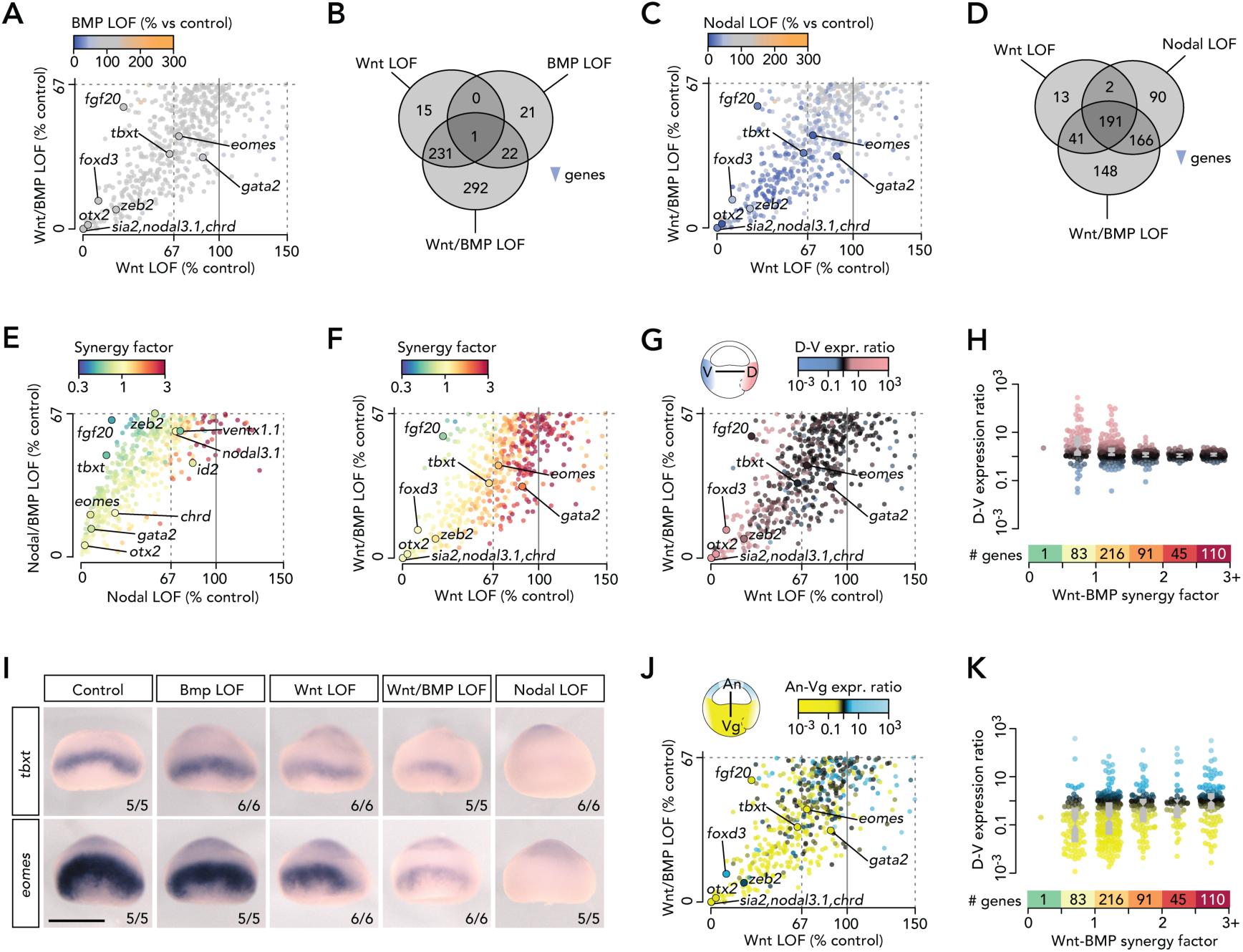
Uniform Gene Expression Across Dorso-Ventral Axis Achieved by Wnt/BMP Synergy. (A,C,E,F,G,J) Scatter plots of relative (% to control) transcript levels between indicated LOFs with each dot (gene) color-coded according to a third attribute: (A,B) relative (% to control) transcript levels, (E,F) synergy factors between single inductive signals, and (G,J) regional expression ratios between opposite ends of the indicated axis. (B,D) Venn diagram of down-regulated genes by indicated LOFs. (H,K) Box and beeswarm plots of regional expression (as measured along the indicated axes) depending on increased Wnt-BMP synergy. (I) WMISH of *tbxt* and *eomes* transcripts under various LOFs. Control and Nodal LOF pictures are from Gentsch et al. (2018b). See also Figure S3 and Table S3.

Overall, the loss of canonical Wnt, Nodal and/or BMP signaling caused the misregulation of ∼39% (∼22.1% down, ∼2.1% down/up and ∼14.4% up) of genes activated at ZGA (Figure 4A). These signals were required for most regional ZGA on the dorsal side (∼89%) and in the VH and MZ (∼82%). Notably, their input affected virtually all genes (∼98%, 56/57) with enriched expression in the dorso-vegetal/MZ quadrant (Figure 4B,C). Thus, Wnt, Nodal and BMP substantially contribute to regional ZGA in most anatomical domains of the early gastrula embryo with the exception of animally enriched transcription (∼19%). Animal- and ventral-specific gene expression relies strongly on both activation by ubiquitous maternal TFs (e.g. Pou5f3/Sox3) and on repression by signals (Figure 4B,D) and other maternal TFs (e.g. VegT) on the opposite side (Figure S2F).

**Figure 4.**
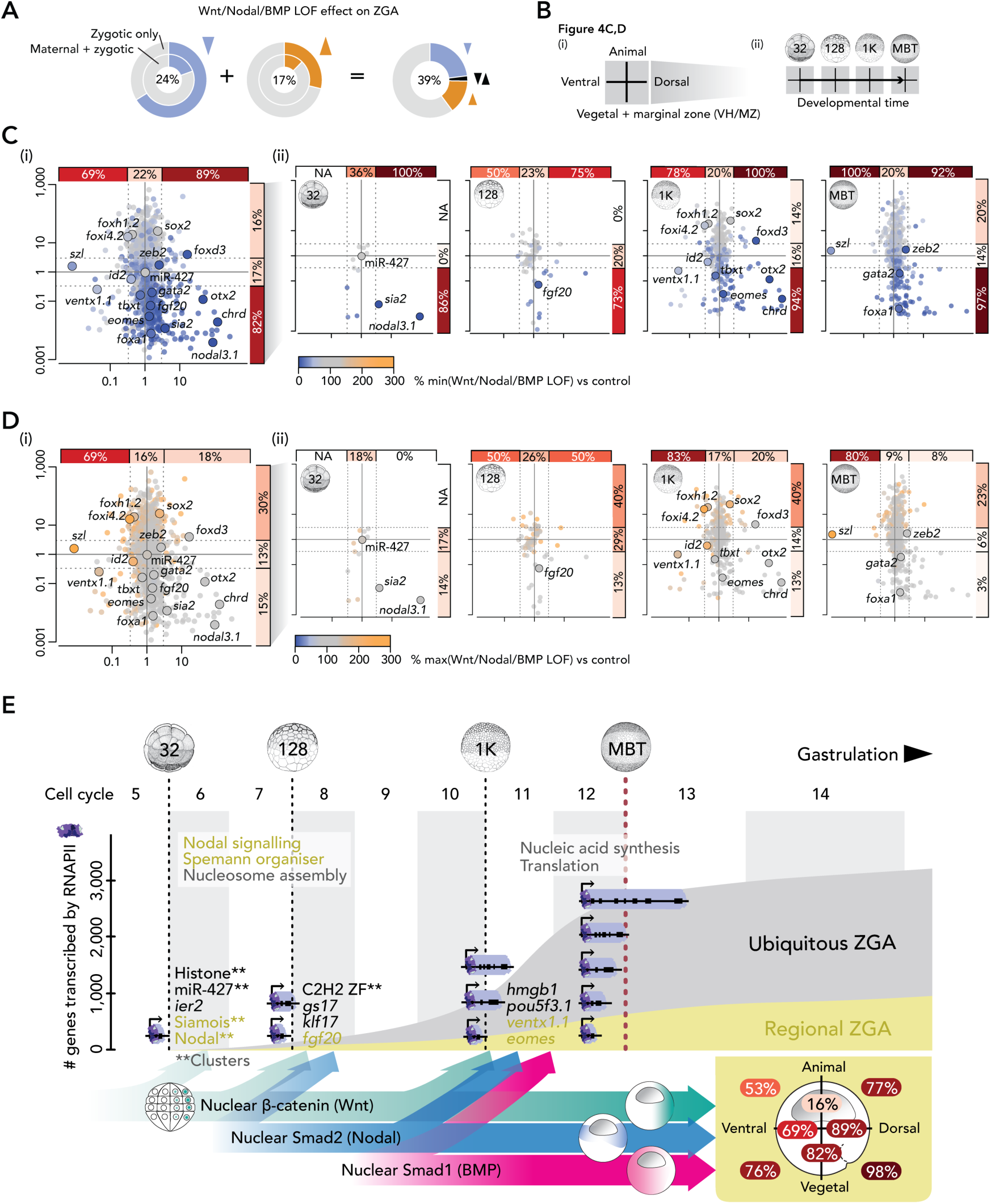
Canonical Wnt, Nodal and BMP Signals Induce the Majority of Regional ZGA. (A) Total percentage of active (zygotic only and maternal-zygotic) genes mis-regulated by Wnt, Nodal and/or BMP LOF. (B) Graphical explanations of figure panels (C) and (D). (C,D) Summary (i) and temporal resolution (ii) of minimal (C) and maximal (D) transcript levels of active genes (separated by regional expression along the primary body axes) detected among Wnt, Nodal and/or BMP LOFs. Percentages only refer to the down-regulated (C) or up-regulated (D) genes among all zygotic genes with the same range of expression ratios along the animal-vegetal or dorso-ventral axes. (E) Exponential activation of gradually longer genes before the MBT (bulk ZGA) when cell divisions occur at rapid and nearly constant intervals (∼20 min at 28°C). Sequential induction of the canonical Wnt, Nodal and BMP pathway is critical to high percentages of regional ZGA (as measured along the two primary body axes within the indicated halves and quadrants of an early gastrula embryo): e.g. ∼89% or ∼98% of gene expression enriched in the dorsal half or vegetal-dorsal quadrant, respectively.

## DISCUSSION

Our study provides two major insights into the mechanisms by which ZGA is initiated in time and space in *Xenopus tropicalis*. The first concerns the temporal aspects, where we find that RNAPII can be detected across gene bodies well before the MBT, during the period when rapid synchronous cell divisions divide the zygote into 4,096 blastomeres (Figure 4E). The average length of genes covered by RNAPII grows during this time, from ∼1 kb at the 32-cell stage to ∼6 kb at the 1,024-cell stage; these figures compare with an average size of ∼16 kb for maternally-expressed genes and for genes expressed after the MBT. Recent long-read sequencing of the zebrafish transcriptome at pre-MBT stages identified transcripts as long as 8 kb spanning multiple pri-miR-430 elements (Nudelman et al., 2018). Furthermore, RNAPII elongation in pre-MBT *Drosophila* embryos occurred at rates of 2.4 to 3.0 kb/min (Chen et al., 2013; Fukaya et al., 2017).

We do not know why RNAPII, despite its high abundance and its ability to promote rapid elongation, is restricted at early stages from transcribing more genes and longer genes. It may, perhaps, be a consequence of the gradual nature of the chromatin remodeling that occurs during these stages, from the accessibility of *cis*-regulatory elements (Gentsch et al., 2018b; Liu et al., 2018; Lu et al., 2016; Wu et al., 2016) to the spatial organisation of an initially unstructured or highly variable chromatin landscape (Du et al., 2017; Flyamer et al., 2017; Hug et al., 2017; Kaaij et al., 2018; Ke et al., 2017). Whatever the reason, the increase of elongated RNAPII engagement between the 128-cell stage and the MBT indicates that a significant component of ZGA disregards the N/C ratio which was originally thought to underlie the onset of transcription at the MBT (Newport and Kirschner, 1982b). Similar conclusions have been drawn from profiling the zygotic transcriptome of haploid *Drosophila* (Lu et al., 2009) and cell cycle-arrested zebrafish (Chan et al., 2018). Thus, it is becoming clear from work in flies, fish and frogs that ZGA starts before the MBT and accelerates thereafter (Ali-Murthy et al., 2013; Collart et al., 2014; Mathavan et al., 2005; Owens et al., 2017; Tan et al., 2013), reaching a peak at the MBT (reviewed by Jukam et al., 2017 and Langley et al., 2014).

These observations notwithstanding, it remains possible that cell cycles do contribute to the temporal progression of ZGA and the exponential increase in the number of activated genes before the MBT. In particular, cell cycles may accelerate chromatin remodeling by displacing suppressors in mitotic chromatin and providing unique access to TFs (Halley-Stott et al., 2014) and structural proteins of high-order chromatin (Ke et al., 2017). For example, maternal core histones have been shown to prevent premature ZGA by competing with specific TFs (Joseph et al., 2017).

In addition to the small sizes of the earliest activated genes, we observed that most of these genes, which have no or few introns, code for groups of related factors like histones or zinc finger TFs, and that they appear as clusters spanning up to several hundred kilobases. This is in line with previous findings of the earliest active multicopy and intron-poor genes like miR-427 and *nodal5/6* in *Xenopus* embryos (Collart et al., 2014; Lund et al., 2009; Owens et al., 2016; Skirkanich et al., 2011; Takahashi et al., 2006; Yanai et al., 2011; Yang et al. 2002) and miR-430 in zebrafish and Medaka fish (Giraldez et al., 2005; Heyn et al., 2014; Tani et al., 2010). The number and spatial proximity of clustered genes enhances transcriptional output by allowing the sharing of multiple cis-regulatory elements (arranged as super-enhancers) (Whyte et al., 2013) and by fortifying transcriptional condensates of TFs, coactivators and RNAPII (Boija et al., 2018; Cho et al., 2018; Chong et al., 2018; Sabari et al., 2018; Shrinivas et al., 2018). Overall, based on enriched gene functions, we discovered that ZGA exerts a temporal control of gene expression from nucleosome remodeling before the MBT to protein degradation after the MBT.

Our second insight concerns spatial ZGA, and the observation that we can assign a large proportion of spatio-temporal ZGA to key signaling pathways (reviewed by Arnold and Robertson, 2009 and Kimelman, 2006). Canonical Wnt, Nodal and BMP signaling govern regional ZGA in line with the nuclear translocation of their signal mediators (Faure et al., 2000; Larabell et al., 1997; Schohl and Fagotto, 2002). Thus, Nodal signaling predominantly affects transcription within the vegetal hemisphere and marginal zone, while Wnt and BMP initiate transcription in dorsal and ventral regions, respectively. The timing of regional ZGA is defined by the sequential translocation of signal mediators such that nuclear β-catenin directs regional ZGA at the 32-cell stage, followed by nuclear Smad2 at the 128-cell stage and Smad1 at the 1,024-cell stage. While Smad2-mediated signal transduction depends on the zygotic transcription of its six Nodal ligands (Faure et al., 2000; Gentsch et al., 2018b; Jones et al., 1995; Yang et al., 2002), canonical Wnt and BMP signaling are initiated by the maternally inherited ligands Wnt11 and BMP2/4/7, respectively (Faure et al., 2000; Heasman, 2006; Tao et al., 2005).

We also show that the synergy of opposing signals of the Wnt and BMP pathway affects many Nodal-dependent genes with uniform expression along the dorso-ventral axis such as *eomes* and *Brachyury* (*tbxt*). It is not yet clear whether Wnt/BMP synergy arises from joint chromatin engagement or from mutual or post-translational interactions. For instance, Wnt8a signal can enhance BMP transcriptional readouts by inhibiting the phosphorylation of GSK3, which normally targets Smad1 for degradation (Fuentealba et al., 2007). However, the analysis of *Brachyury* gene regulation in zebrafish suggests that Wnt and BMP can be integrated at a single cis-regulatory DNA element and together with a separate Nodal-responsive DNA element they can establish uniform dorso-ventral expression (Harvey et al., 2010). This is further corroborated by our analysis of genome-wide chromatin engagement (Gentsch et al., 2018b): the canonical DNA recognition motif for the Wnt-associated basic helix-span-helix (bHSH) TF AP-2 was more enriched at Smad1 than Smad2 binding sites (Figure S3O).

We therefore propose that Wnt, BMP and Nodal signal mediators are critical to regional ZGA and that they balance initially opposing cell fate commitments. However, we have previously shown that signal integration also relies on maternal pioneer TFs like Pou5f3 and Sox3 to make signal-responsive *cis*-regulatory elements accessible for signal mediator binding. For example, Nodal-induced transcription of the *Brachyury* gene depends on the pioneering roles of maternal Pou5f3 and Sox3, and less on their transcriptional activities (Gentsch et al., 2018b).

Overall, we demonstrate that the temporal and spatial dynamics of regional ZGA are regulated by the sequential and spatially restricted translocation of Wnt, Nodal and BMP signal mediators. These events establish the formation of the primary body axes and germ layers of the embryo. Temporal RNAPII profiling indicates that >1,000 genes of increasing length are activated before MBT and that this substantial portion of ZGA is independent of both the classic N/C ratio and of cell cycle lengthening.

## LIMITATIONS OF THE STUDY

We detected a dramatic increase in genome-wide recruitment of RNAPII over the cleavage stages during which the genome begins to be transcribed. We used within-sample normalization to scale developmental stage-specific RNAPII profiles. However, because of the large differences in total RNAPII enrichment between samples, chromatin spike-ins are now considered a more accurate method to normalize ChIP-Seq profiles across consecutive developmental stages (Chen et al., 2016). We combined separate whole-embryo determinations of RNAPII engagement and transcript levels to reveal the temporal dynamics of ZGA. This approach could be improved by profiling RNAPII-associated RNA to directly couple RNAPII elongation with transcript accumulation (e.g., Churchman and Weissman, 2011). In addition, the spatial resolution of ZGA, which is based on transcriptomics of dissected embryonic parts in our study, could be enhanced by various deep single-cell profiling and super-resolution imaging technologies. We show that most regional ZGA depends on Wnt, Nodal and BMP signals, but an important question remains: How are these signals integrated at the chromatin level to sustain RNAPII-mediated transcript elongation? In part, this could be investigated by targeted genome editing to increase our understanding of signal-responsive gene regulatory DNA.

## ACKNOWLEDGEMENTS

We thank Abdul Sesay, Leena Bhaw, Harsha Jani, Deborah Jackson and Meena Anissi for deep sequencing; Mareike Thompson for critical reading of the manuscript; and the Smith lab for discussions and advice. G.E.G and J.C.S. were supported by the Medical Research Council (program number U117597140) and are now supported by the Francis Crick Institute, which receives its core funding from Cancer Research UK (FC001-157), the UK Medical Research Council (FC001-157), and the Wellcome Trust (FC001-157).

## AUTHOR CONTRIBUTIONS

Conceptualization, G.E.G.; Methodology, G.E.G.; Computational Code, G.E.G.; Formal Analysis, G.E.G. and N.D.L.O.; Investigation, G.E.G.; Writing – Original Draft, G.E.G. and J.C.S.; Writing – Review & Editing, G.E.G and J.C.S.; Funding Acquisition, J.C.S.

## SUPPLEMENTAL INFORMATION

**Figure S1.**
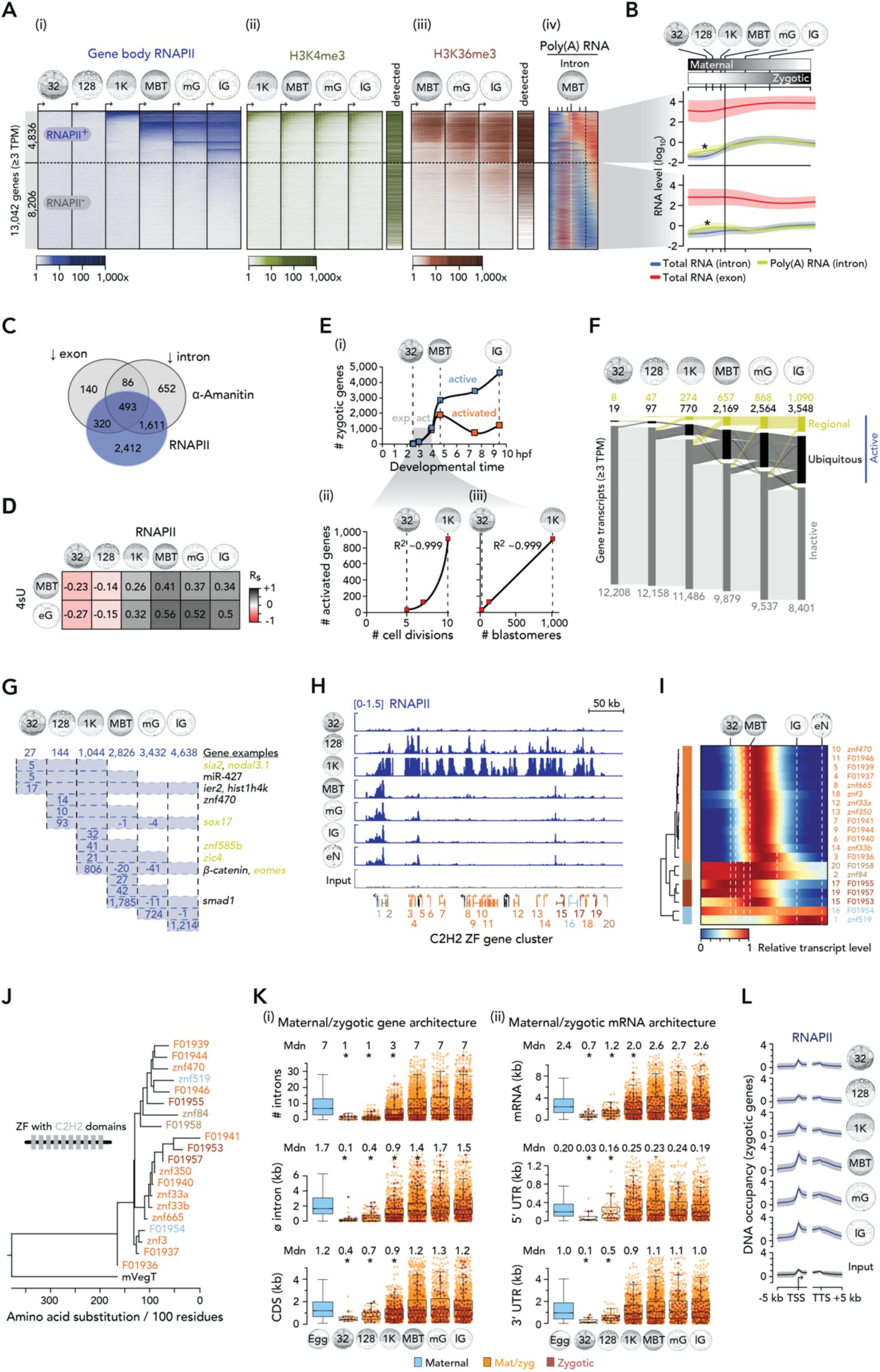
Dynamics and Architecture of ZGA in *X. tropicalis*, Related to Figure 1. (A) Progression of ZGA from the 32-cell to the late gastrula stage based on (i) whole gene body (full-length) occupancy of RNAPII (i.e., RNAPII was enriched across entire gene bodies; see Transparent Methods). Co-aligned: Active histone marks H3K4me1 (ii) and H3K36me3 (iii) (Hontelez et al., 2015) and intronic signal from the high time-resolution profile of poly(A) RNA (iv) (Owens et al., 2016). The horizontal dotted line separates RNAPII-engaged (RNAPII^+^) from non-engaged (RNAPII^−^) genes as detected until the late gastrula stage. The vertical dotted lines in the poly(A) RNA plot indicate the developmental time points of the MBT (white) and the late gastrula stage (black), respectively. (B) Transcript feature levels (mean +/- SD) during the maternal-to-zygotic transition. Asterisk, polyadenylation immediately after fertilization (Collart et al., 2014) transiently increased the intronic signal obtained from the poly(A) RNA samples. (C) Venn diagram of zygotic genes detected by full-length RNAPII occupancy or reduced exonic or intronic transcript counts upon blocking RNAPII-mediated transcription with α-amanitin. (D) Pairwise Spearman’s correlations (R_s_) of enrichment values resulting from RNAPII profiling and 4sU tagging to detect zygotic genes at indicated developmental stages. (E) Plots of the number of active and newly activated genes (i) or newly activated genes versus the developmental time (i), the number of completed cell divisions (ii) or formed blastomeres (iii). (F) Alluvial diagram of spatio-temporal ZGA including maternally inherited RNA transcripts of genes not activated by the late gastrula stage. Tissue-specificity inferred from regional transcript enrichments along the animal-vegetal or the dorso-ventral or both axes (Blitz et al., 2017). (G) Numbers of genes with full-length RNAPII occupancy at indicated developmental stages. Examples of ubiquitously (black) and tissue-specifically (orange) expressed genes are listed to the right. (H) RNAPII dynamics at the Cys2-His2 [C2H2] zinc finger (ZF) cluster from the 32-cell to the early neurula stage. (J) Expression dynamics of C2H2 ZF genes normalized to maximal transcript levels recorded between fertilization and 23.5 hpf (Owens et al., 2016). (J) Phylogenetic tree of the C2H2 ZF genes shown in (H,I). Maternal VegT (mVegT), outgroup TF of this phylogenetic tree. (K) Box and beeswarm plots showing various metrics of the zygotic/maternal genes (i) and mRNA (ii) during ZGA. Asterisks, significant Wilcoxon rank-sum tests against maternal and post-MBT activated genes and corresponding effect sizes (r_effect_): # introns, p <2.1e-13, r_effect_ 0.07-0.43; Ø intron (kb), p <7.9e-7, r_effect_ 0.07-0.42; CDS (kb), p <3e-7, r_effect_ 0.05-0.27; mRNA (kb), p <7e-11, r_effect_ 0.06-0.32; 5’ UTR (kb), p <0.015, r_effect_ 0.03-0.15; and 3’ UTR (kb), p <1.4e-6, r_effect_ 0.04-0.21. (L) Meta-profiles (mean +/- SD) of RNAPII (separated by developmental stage) and input (negative control) densities at zygotic genes. Abbreviations 32, 32-cell; 128, 128-cell; 1K, 1,024-cell; MBT, mid-blastula transition; mG, mid-gastrula; lG, late gastrula; eN, early neurula; 4sU, 4-thiouridine; Mdn, median; TPM, transcripts per million.

**Figure S2.**
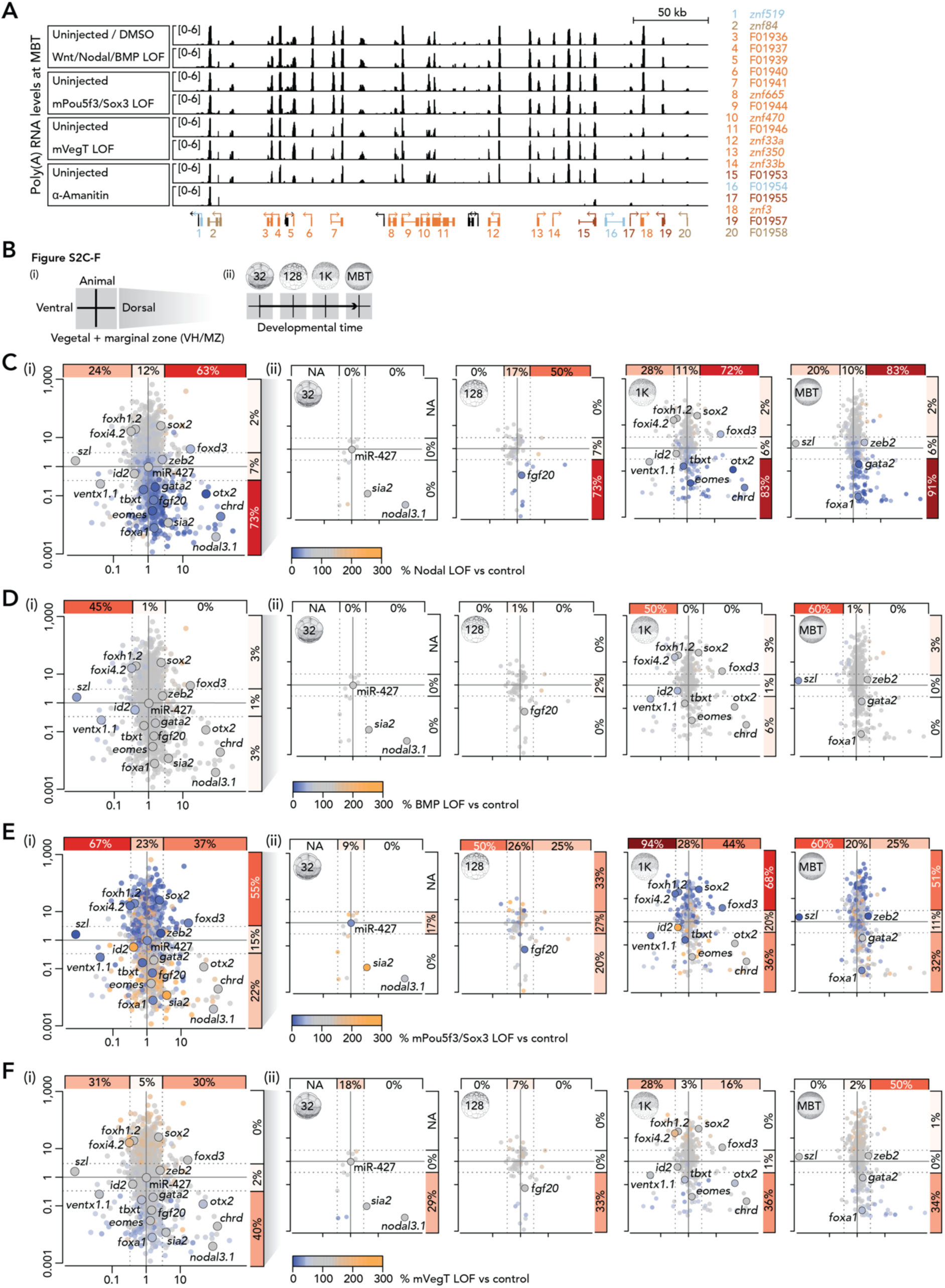
Effect of Canonical Wnt, Nodal and BMP Signals on ZGA, Related to Figure 2. (A) Poly(A) RNA profiles of the C2H2 ZF cluster (Figure S1H) for indicated control and LOFs at the MBT. (B) Graphical explanations of figure panels (C-F). (C-F) Summary (i) and temporal resolution (ii) of gene mis-regulations upon the LOF of Nodal (C) or BMP (D) signaling or maternal Pou5f3/Sox3 (mPou5f3/Sox3) (E) or VegT (mVegT) (F). Percentages only refer to the down-regulated genes (by ≥1/3 compared to control expression level) among all zygotic genes with the same range of expression ratios along the animal-vegetal or the dorso-ventral axes.

**Figure S3.**
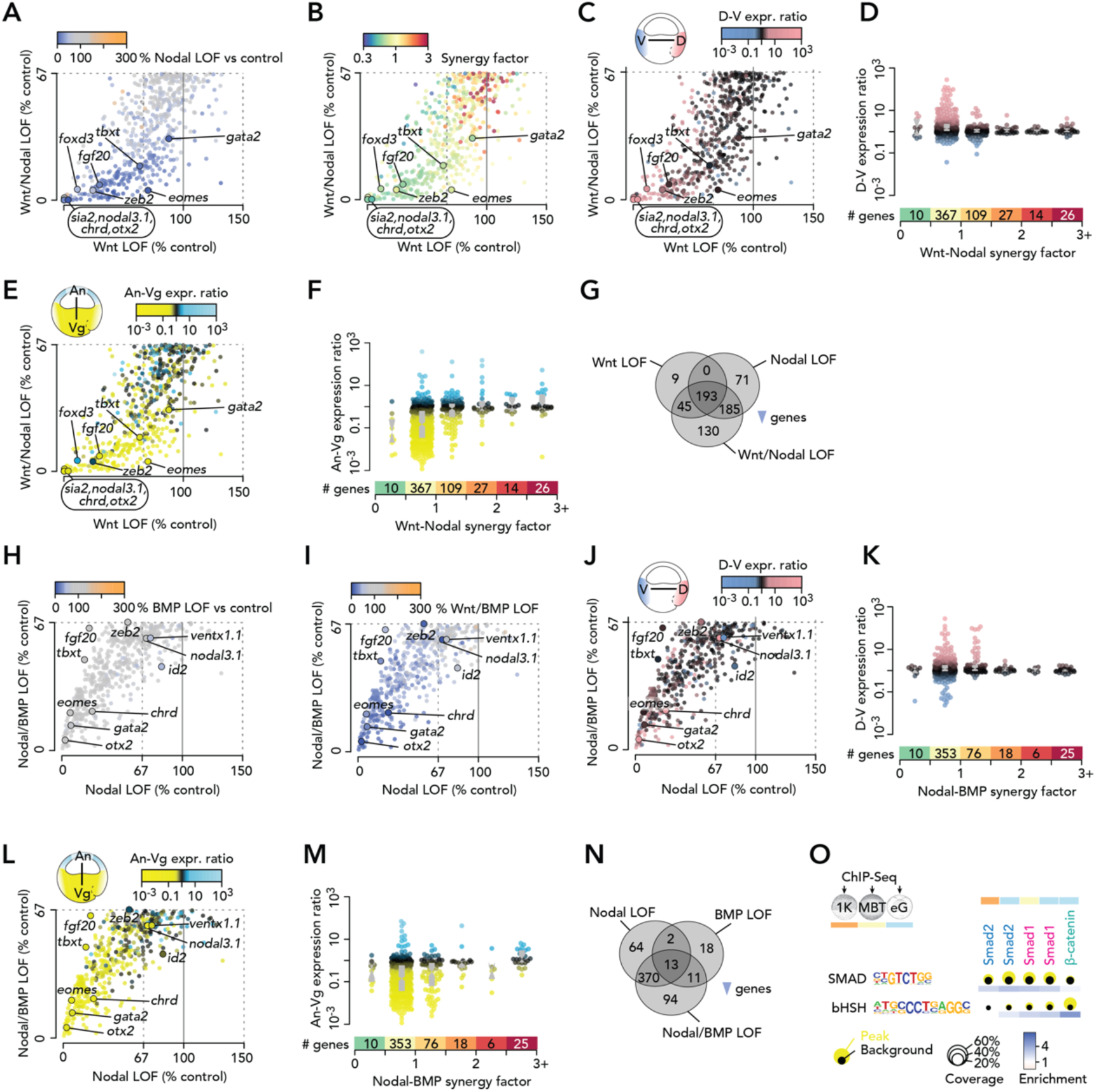
Relationship between Canonical Wnt, Nodal and BMP to Control Regional ZGA, Related to Figure 3. (A-C,E,H-J,L) Scatter plots of relative (% to control) transcript levels between indicated LOFs with each dot (gene) color-coded according to a third attribute: (A,H,I) relative (% to control) transcript levels, (B) synergy factors between single inductive signals, and (C,E,J,L) regional expression ratios between opposite ends of the indicated axis. (G,N) Venn diagram of down-regulated genes by indicated LOFs. (D,F,K,M) Box and beeswarm plots of regional expression (as measured along the indicated axes) depending on increased Wnt-Nodal (D,F) or Nodal-BMP (K,M) synergy. (O) Coverage and enrichment of Smad and β-catenin-associated DNA motifs (SMAD and bHSH motifs) at endogenous binding sites of β-catenin, Smad1, Smad2 (Gentsch et al., 2018b) at indicated developmental stages (color-coded).

## SUPPLEMENTAL TABLES

**Table S1.**
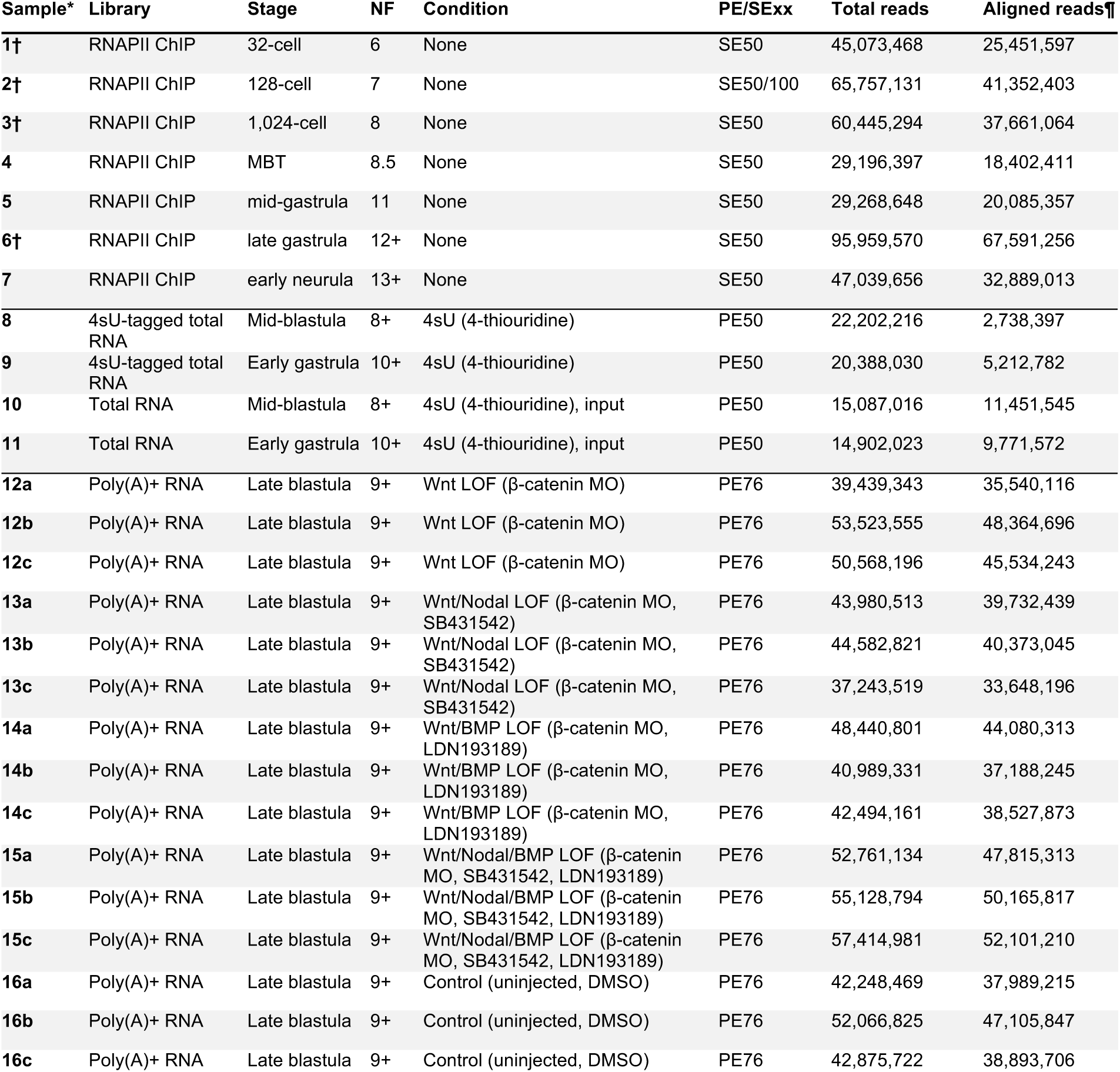
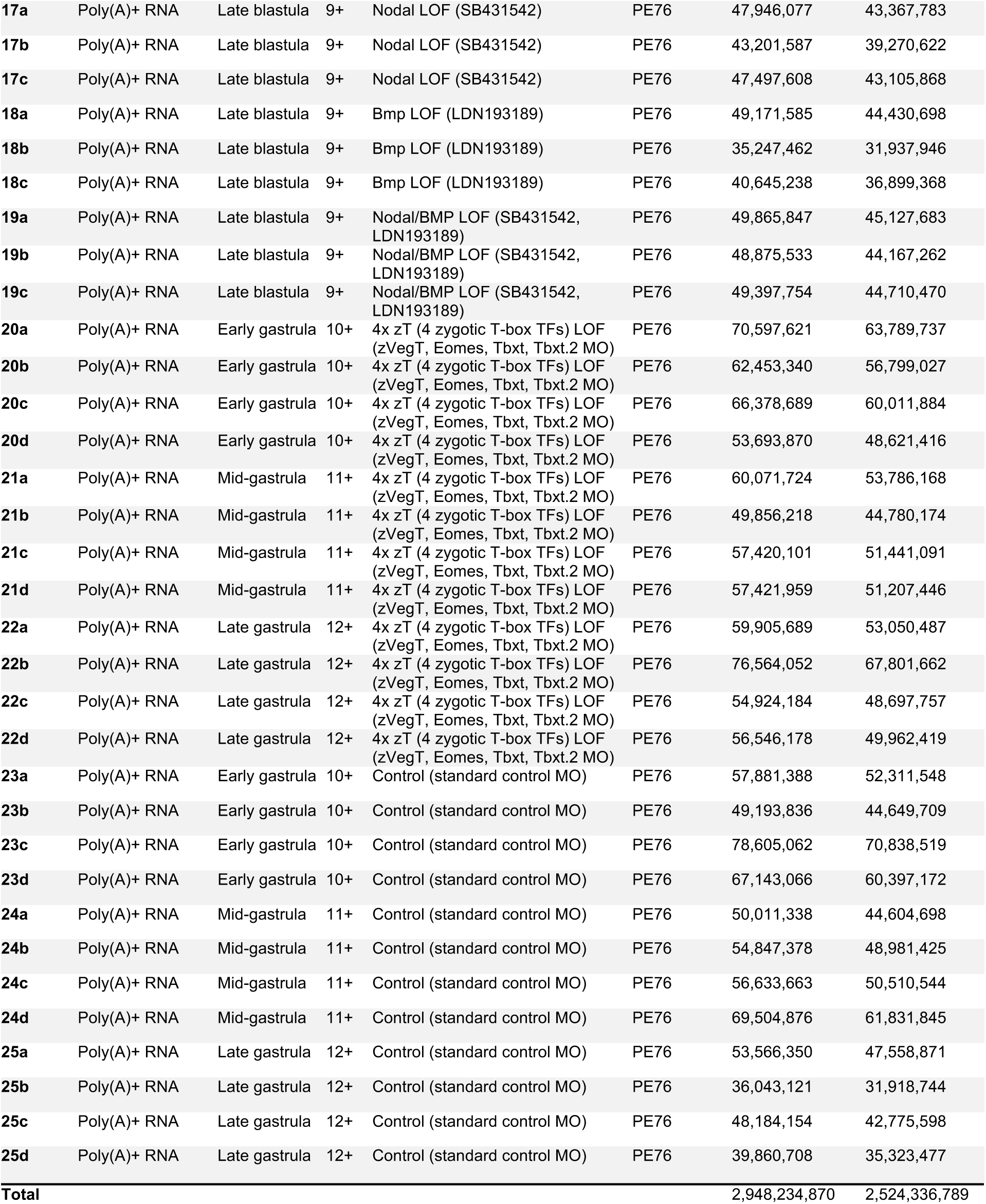
Summary of Deep sequencing and Read Alignments, Related to Figure 1 and 2. The meta-data of ChIP-Seq, 4sU-Seq and RNA-Seq runs includes the developmental stage, condition, read type, and total and genome-aligned read numbers. *, all rows represent manually collected biological sample: a, b and c mark biological replicates. †, library was sequenced twice. Reads were pooled from both sequencing runs. ¶, reported numbers are reads that are non-redundantly aligned to the genome assembly 7.1 with a mapping quality (MAPQ) of 10 (DNA) and 255 (RNA). NF, staging according to Nieuwkoop and Faber (1994).

**Table S2. Temporal Progression of ZGA, Related to Figure 1.**

Activated genes are listed according to the earliest developmental stage of full-length RNAPII occupancy. The lists also include the overall enrichment of RNAPII across the gene body, the maternal contribution (TPM) (Owens et al., 2016) and the enrichment of de novo synthesized transcripts determined by 4sU tagging at the MBT and the early gastrula stage.

**Table S3. Differential ZGA Analysis, Related to Figure 2, 3 and 4.**

Differential expression analysis of genes showing ≥50% (FDR ≤10%) reduced transcript levels in α-amanitin-injected embryos. Normalized transcript levels (inferred from exon or intron counts) are scaled (percentage, %) to the expression level in control embryos. The list also includes FDRs, expression ratios across the animal-vegetal or dorso-ventral axis (Blitz et al., 2017), the earliest developmental stage of full-length RNAPII occupancy, and average expression levels between 0 and 1 hpf (maternal) and 5 and 9 hpf (from the MBT to the mid-gastrula stage).

**Movie S1. Quadruple LOF of Zygotic T-box TFs, Related to Figure 2.**

Simultaneous filming of the vegetal (top row) and animal (bottom row) hemisphere of 4x zT LOF (labelled as T-box KD in the movie) (left) and control (right) embryos from early gastrula to mid-tailbud stage.

## TRANSPARENT METHODS

### CONTACT FOR REAGENTS AND RESOURCE SHARING

Further information and requests for resources and reagents should be directed to and will be fulfilled by the Lead Contact, James C. Smith.

### EXPERIMENTAL MODEL AND SUBJECT DETAILS

#### Xenopus tropicalis Manipulation

Standard procedures were used for ovulation, fertilization, and manipulation and incubation of embryos (Khokha et al., 2002; Sive et al., 2000). Briefly, frogs were obtained from Nasco (Wisconsin, USA). Ovulation was induced by injecting serum gonadotropin (Intervet) and chorionic gonadotropin (Intervet) into the dorsal lymph sac of mature female frogs. Eggs were fertilized *in vitro* with sperm solution consisting of 90% Leibovitz’s L-15 medium (Thermo Fisher Scientific, Cat#11415064) and 10% fetal bovine serum (Thermo Fisher Scientific, Cat#10500056). After 10 min, fertilized eggs were de-jellied with 2.2% (w/v) L-cysteine (Merck, Cat#168149) equilibrated to pH 8.0. Embryos were cultured in 5% Marc’s Modified Ringer’s solution (MMR) (5 mM NaCl, 0.1 mM KCl, 0.1 mM CaCl_2_, 0.05 mM MgSO_4_ and 0.25 mM HEPES pH7.5) at 21°C-28°C. Embryos were staged according to Nieuwkoop and Faber (1994). All *Xenopus* work fully complied with the UK Animals (Scientific Procedures) Act 1986 as implemented by the Francis Crick Institute.

#### Chromatin immunoprecipitation (ChIP)

ChIP was carried out as detailed previously (Gentsch and Smith, 2017). Briefly, de-jellied *X. tropicalis* embryos were fixed at room temperature with 1% formaldehyde (Merck, Cat#F8775) in 1% MMR for 25 min. The fixation time was extended to 45 min for pre-gastrula stages. The following number of embryos were used for ChIP-Seq: 1,400 at the 32-cell stage, 1,000 at the 128-cell stage, 700 at the 1,024-cell stage, 450 at the MBT and 350 for the post-MBT stages. Fixation was terminated by rinsing embryos three times with ice-cold 1% MMR. Fixed embryos were homogenized in CEWB1 (150 mM NaCl, 1 mM EDTA, 1% (v/v) Igepal CA-630 [Merck, Cat#I3021], 0.25% (w/v) sodium deoxycholate [Merck, Cat#SRE0046], 0.1% (w/v) sodium dodecyl sulfate [Merck, Cat#71729] and 10 mM Tris-HCl pH 8.0) supplemented with 0.5 mM DL-Dithiothreitol (Fluorochem, Cat#M02712) and protease inhibitors (Roche, Cat#11873580001). The homogenate was left on ice for 5 min and then centrifuged at 1,000 g (4°C) for 5 min. Homogenization and centrifugation was repeated once before resuspending the pellet in 1-3 ml CEWB1. Chromatin was solubilized and fragmented by microtip-mediated ultra-sonication (Misonix 3000 sonicator with a tapered 1/16-inch microtip). The solution of fragmented chromatin was cleared by centrifuging at 16,000 g (4°C) for 5 min. About 1% of the cleared chromatin extract was set aside for the input sample (negative control). The remaining chromatin was incubated overnight at 4°C on a vertical rotor (10 rpm) with 20 µl of the mouse monoclonal anti-RNAPII (8WG16) (Covance, Cat#MMS-126R; RRID: AB_10013665) antibody. After adding 100 µl of washed protein G magnetic beads (Thermo Fisher Scientific, Cat#10003D) the solution was incubated for another 4 h at 4°C on a vertical rotor (10 rpm). The beads were washed eight times in CEWB1 and once in TEN (10 mM Tris-HCl pH 8.0, 150 mM NaCl and 1 mM EDTA) at 4°C. ChIP was eluted off the beads twice with 100 µl SDS elution buffer (50 mM Tris-HCl pH 8.0, 1 mM EDTA and 1% (w/v) sodium dodecyl sulfate) at 65°C. ChIP eluates were pooled before reversing DNA-protein cross-links. Input (filled up to 200 µl with SDS elution buffer) and ChIP samples were supplemented with 10 µl 5 M NaCl and incubated at 65°C for 6-16 h. Samples were treated with proteinase K (Thermo Fisher Scientific, Cat#AM2548) and RNase A (Thermo Fisher Scientific, Cat#12091021) to remove any proteins and RNA from the co-immunoprecipitated DNA fragments. The DNA was purified with phenol:chloroform:isoamyl alcohol (25:24:1, pH 7.9) (Thermo Fisher Scientific, Cat#AM9730) using 2.0-ml Phase Lock Gel Heavy microcentrifuge tubes (VWR, Cat#733-2478) for phase separation and precipitated with 1/70 volume of 5 M NaCl, 2 volumes of absolute ethanol and 15 µg GlycoBlue (Thermo Fisher Scientific, Cat#AM9516). After centrifugation, the DNA pellet was air-dried and dissolved in 11 µl elution buffer (10 mM Tris-HCl pH 8.5). The DNA concentration was determined on a Qubit fluorometer using high-sensitivity reagents for detecting double-stranded DNA (10 pg/µl to 100 ng/µl) (Thermo Fisher Scientific, Cat#Q33231).

#### ChIP-Seq Library Preparation

Using the KAPA Hyper Prep Kit (Roche, Cat#KK8504), 2.5-5 ng ChIP DNA or 5 ng input DNA were converted into indexed paired-end libraries as previously described (Gentsch and Smith, 2017). Briefly, DNA fragments were end-repaired and A-tailed for 30 min at 20°C followed by 30 min at 65°C before cooling to 4°C. 7.5 pmol TruSeq (single index) Y-adapters (IDT) were ligated to the DNA fragment ends for 20 min at 20°C. The DNA ligation product was extracted with 0.8x SPRI (solid phase reversible immobilisation) beads (Beckman Coulter, Cat#A63882) and amplified in five PCR cycles (15 sec at 98°C, 30 sec at 60°C and 30 sec at 72°C) using the KAPA high-fidelity polymerase master mix (Roche, Cat#KK2602) and 25 pmol Illumina P5 (forward) and P7 (reverse) primers (IDT). After cleaning up the PCR reaction with 1× SPRI beads, the DNA library was size-separated by electrophoresis using E-gel EX agarose gels (Thermo Fisher Scientific, Cat#G401002). A gel slice containing DNA ranging from 250 to 450 bp in size was dissolved shaking in 350 µl QG buffer (Qiagen) using a thermomixer (1,000 rpm) at room temperature. The DNA was purified with MinElute columns (Qiagen, Cat#28604) and eluted off these columns twice using 11 µl elution buffer (10 mM Tris-HCl pH 8.5). The library was re-amplified using another 6-8 PCR cycles yielding 100-200 ng DNA without adapter dimer contamination. The DNA library was cleaned up with 1x SPRI beads.

#### Illumina Sequencing

All sequencing libraries were quality controlled: The DNA yield and fragment size distribution were determined by fluorometry and chip-based capillary electrophoresis, respectively. ChIP-Seq and RNA-Seq libraries were sequenced on the Illumina HiSeq 2500 and 4000, respectively, by the Advanced Sequencing Facility of the Francis Crick Institute. Sequencing samples and read alignment results are summarized in Table S1.

#### Post-Sequencing Analysis of ChIP-Seq

Single reads of maximal 50 bases were processed using trim_galore v0.4.2 (Babraham Institute, UK) to trim off low-quality bases (default Phred score of 20, i.e. error probability was 0.01) and adapter contamination from the 3’ end. Processed reads were aligned to the *X. tropicalis* genome assembly v7.1 and v9.1 (for Hilbert curves) running Bowtie2 v2.2.9 (Langmead and Salzberg, 2012) with default settings (Table S1). Alignments were converted to the HOMER’s tag density format (Heinz et al., 2010) with redundant reads being removed (makeTagDirectory -single -tbp 1 -unique -mapq 10 -fragLength 175 -totalReads all). Only uniquely aligned reads (i.e. MAPQ ≥10) were processed. We pooled all input alignments from various developmental stages (Gentsch et al., 2018b). This created a comprehensive mappability profile that covered ∼400 million unique base pair positions. For Hilbert curves, tag densities were generated across the genome v9.1 using sliding (200-bp increments) 400-bp window. Background signals (<0.3 reads per 1 million mapped reads) were removed. Blacklisted (Gentsch et al., 2018b) regions (except for MIR-427) were excluded using intersectBed (-v -f 0.5) from BEDtools v2.25.0 (Quinlan and Hall, 2010).

#### Detecting Zygotic and Maternal Genes Using RNAPII Profiling and High Time-Resolution Transcriptomics

Normalized RNAPII and input tag densities were calculated across the gene body in 10 bins of equal size. Gene annotations v7.1 were altered based on a few known zygotic isoforms and some corrections obtained from assembling total and poly(A) RNA (Owens et al., 2016) from stage 6 to stage 12.5 *de novo* (Pertea et al., 2016). A few genes had previously been annotated as gene clusters due to assembly uncertainties. We reduced the annotation of polycistronic MIR-427 to the minus arm (scaffold_3b:3516900-3523400) and only monitored *nodal3.5* and *nodal5.3* within their respective gene clusters. Gene bodies with <40% mappability were removed. Here, the threshold of mappability per bin was set at 10% of the input read density averaged across all gene bodies in use. Subsequently, enrichment values were only obtained for all mappable bins by dividing read densities of RNAPII and input. Further, we restricted the analysis to genes for which ≥3 transcripts per million (TPM) could be detected on average over three consecutive time points (i.e. over the developmental time of 1 h) of a high-resolution profile of total RNA (Owens et al., 2016) from fertilization to after gastrulation (stage 13). Genes were considered active when RNAPII enrichments along their full length (see thresholds below) and corresponding transcripts (≥0.1 TPM) were simultaneously detected. Transcript levels were calculated over three consecutive time points +/- 1 h from the developmental stage of RNAPII profiling. RNAPII enrichment covered ≥80% of the mappable gene body and reached at least one of the following thresholds: (1) 2.6-fold, (2) 1.8-fold and 1.4-fold at the next or previous stage, (3) 1.4-fold and 1.8-fold at the next or previous stage, or (4) 1.4-fold over three consecutive stages. The heatmap (Figure 1B and S1A) was sorted by the developmental stage (1^st^) and the overall fold (2^nd^) of RNAPII enrichment. Zygotic and maternal contributions to transcriptome (Figure 1G) were based on RNAPII enrichment (see above) and mean transcript levels (≥0.1 TPM) detected between 0 and 1 hpf, respectively.

#### Peak Calling and Motif Enrichment Analysis

Peak calling and motif enrichment analysis were carried out as previously reported (Gentsch et al., 2018b). Briefly, HOMER v4.8.3 (Heinz et al., 2010) was used to identify the binding sites of Smad1 (Gentsch et al., 2018b), Smad2 (Chiu et al., 2014; Gentsch et al., 2018b; Yoon et al., 2011) and β-catenin (Gentsch et al., 2018b; Nakamura et al., 2016) by virtue of ChIP-enriched read alignments (hereafter called peaks): findpeaks -style factor -minDist 175 -fragLength 175 - inputFragLength 175 -fdr 0.001 -gsize 1.435e9 -F 3 -L 1 -C 0.97. This means that both ChIP and input alignments were extended 3’ to 175 bp for the detection of significant (FDR ≤0.1%) peaks being separated by ≥175 bp. The effective size of the *X. tropicalis* genome assembly v7.1 was set to 1.435 billion bp, an estimate obtained from the mappability profile (Gentsch et al., 2018b). These peaks showed equal or higher tag density than the surrounding 10 kb, ≥3-fold more tags than the input and ≥0.97 unique tag positions relative to the expected number of tags. To further eliminate any false positive peaks, we removed any peaks with <0.5 CPM and those falling into blacklisted regions showing equivocal mappability due to genome assembly errors, gaps or simple/tandem repeats. Regions of equivocal mappability were identified by a two-fold lower (poor) or three-fold higher (excessive) read coverage than the average detected in 400-bp windows sliding at 200-bp intervals through normalized ChIP input and DNase-digested naked genomic DNA (Gentsch et al., 2018b). All identified regions ≤800 bp apart were subsequently merged. Gap coordinates were obtained from the Francis Crick mirror site of the UCSC genome browser (http://genomes.crick.ac.uk). Simple repeats were masked with RepeatMasker v4.0.6 (Smit et al.) using the crossmatch search engine v1.090518 (Phil Green) and the following settings: RepeatMasker -species “xenopus silurana tropicalis” -s -xsmall. Tandem repeats were masked with Jim Kent’s trfBig wrapper script of the Tandem Repeat Finder v4.09 (Benson, 1999) using the following settings: weight for match, 2; weight for mismatch, 7; delta, 7; matching probability, 80; indel probability, 10; minimal alignment score, 50; maximum period size, 2,000; and longest tandem repeat array (-l), 2 [million bp]. The enrichment and occurrence of predetermined DNA binding motifs was calculated using 100 bp centred across the top 2,000 peaks per chromatin feature and developmental stage: findMotifsGenome.pl -size 100 -mknown -nomotif.

#### Injections and Treatments of Embryos

Microinjections were performed using calibrated needles and embryos equilibrated in 4% (w/v) Ficoll PM-400 (Merck, Cat#F4375) in 5% MMR. Microinjection needles were generated from borosilicate glass capillaries (Harvard Apparatus, GC120-15) using the micropipette puller Sutter p97. Maximally three nanolitres were injected into the animal hemisphere of de-jellied zygotes using the microinjector Narishige IM-300. Embryos were transferred to fresh 5% MMR (without Ficoll PM-400) once they reached about the mid-blastula stage.

For profiling the nascent transcriptome, embryos were injected with 75 ng 4-thiouridine-5’-triphosphate (4sU) (TriLink BioTechnologies, Cat#N-1025), which is incorporated into newly synthesized transcripts.

Loss-of-functions (LOFs) were generated by treating embryos with small molecule inhibitors and/or injecting them with morpholinos (MOs) or α-amanitin. MOs were designed and produced by Gene Tools (Oregon, USA) to block splicing (MO_splice_) or translation (MO_transl_): maternal Pou5f3/Sox3 (mPou5f3/Sox3) LOF, 5 ng *Pou5f3.2* MO_transl_ (Chiu et al., 2014; Gentsch et al., 2018b; GCTGTTGGCTGTACATAGTGTC), 5 ng *Pou5f3.3* MO_transl_ (Chiu et al., 2014; Gentsch et al., 2018b; TACATTGGGTGCAGGGACCCTCTCA) and 5 ng *Sox3* MO_transl_ (Gentsch et al., 2018b; GTCTGTGTCCAACATGCTATACATC); maternal VegT (mVegT) LOF, 10 ng m*VegT* MO_transl_ (Gentsch et al., 2018b; Rana et al., 2006; TGTGTTCCTGACAGCAGTTTCTCAT); canonical Wnt LOF, 5 ng *β-catenin* MO_transl_ (Heasman et al., 2000; TTTCAACAGTTTCCAAAGAACCAGG); LOF of four zygotic T-box TFs (4x zT LOF), 2.5 ng *tbxt (Xbra, t)* MO_splice_ (Gentsch et al., 2013; TGGAGAGACCCTGATCTTACCTTCC), 2.5 ng *tbxt (Xbra, t)* MO_transl_ (Gentsch et al., 2013; GGCTTCCAAGCGCACACACTGGG), 2.5 ng *tbxt.2 (Xbra3, t2)* MO_splice_ (Gentsch et al., 2013; GAAAGGTCCATATTCTCTTACCTTC), 2.5 ng *tbxt.2* (*Xbra3, t2*) MO_transl_ (Gentsch et al., 2013; AGCTGTGCCTGTGCTCATTGTATTG), 5 ng z*VegT* MO_transl_ (Fukuda et al., 2010; Gentsch et al., 2013; CATCCGGCAGAGAGTGCATGTTCCT) and 5 ng *eomes* MO_splice_ (Fukuda et al., 2010; Gentsch et al., 2013; GAACATCCTCCTGCAAAGCAAAGAC); control MO, 5-20 ng standard control MO (CCTCTTACCTCAGTTACAATTTATA) according to the dose used for the β-catenin, mVegT and 4x zT LOF experiment; and 30 pg α-amanitin (BioChemica, Cat#A14850001). To block Nodal (Nodal LOF) and BMP (BMP LOF) signaling, embryos were treated with 100 μM SB431542 (Tocris, Cat#1614) and/or 10 μM LDN193189 (Selleckchem, Cat#S2618) from the 8-cell stage onwards. Control embryos were treated accordingly with DMSO, in which these antagonists were dissolved. Transcriptional effects of combinatorial signal LOF were determined at late blastula stage (stage 9^+^), while those of all other maternal LOFs were determined over three consecutive time points: the MBT (stage 8+), the late blastula (stage 9^+^) and the early gastrula (stage 10^+^) stage. The 4x zT LOF was transcriptionally profiled at early, mid and late gastrula stage (stage 10^+^, 11^+^ and 12^+^). The 4x zT LOF comparison has four biological replicates (n=4). All other comparisons entail three biological replicates (n=3).

#### Extraction of Total RNA

Embryos were homogenized in 800 µl TRIzol reagent (Thermo Fisher Scientific, Cat#15596018) by vortexing. The homogenate was either snap-frozen in liquid nitrogen and stored at −80°C or processed immediately. For phase separation, the homogenate together with 0.2x volume of chloroform was transferred to pre-spun 2.0-ml Phase Lock Gel Heavy microcentrifuge tubes (VWR), shaken vigorously for 15 sec, left on the bench for 2 min and spun at ∼16,000 g (4°C) for 5 min. The upper phase was mixed well with one volume of 95- 100% ethanol and spun through the columns of the RNA Clean & Concentrator 25 Kit (Zymo Research, Cat#R1017) at ∼12,000 g for 30 sec. Next, the manufacturer’s instructions were followed for the recovery of total RNA (>17 nt) with minor modifications. First, the flow-through of the first spin was re-applied to the column. Second, the RNA was treated in-column with 3 U Turbo DNase (Thermo Fisher Scientific, Cat#AM2238). Third, the RNA was eluted twice with 25 µl molecular-grade water. The concentration was determined on the NanoDrop 1000 spectrophotometer or by fluorometry before depleting ribosomal RNA from total RNA (Profiling the Nascent Transcriptome).

#### Tagging the Nascent Transcriptome

Thirty 4sU-injected embryos were collected at the MBT and the early-to-mid gastrula stage. Total RNA was extracted as outlined above. The 4sU-tagging was performed according to Gay et al. (2014) with few minor modifications. The RNA Clean & Concentrator 5 Kit (Zymo Research, Cat#R1013) was used to purify RNA. Briefly, the Ribo-Zero Gold rRNA Removal Kit (Illumina, Cat#MRZG126) was used according to the manufacturer’s instructions to deplete ribosomal RNA from ∼10 µg total RNA. The RNA was purified and fragmented for 4 min at 95°C using the NEBNext Magnesium RNA Fragmentation Module (NEB, Cat#E6150). The RNA was purified again before conjugating HDPD-Biotin (Thermo Fisher Scientific, Cat#21341) to 4sU via disulfide bonds for 3 h in the dark. Purified RNA was mixed with Streptavidin beads (Thermo Fisher Scientific, Cat#65305) to pull down biotin-tagged RNA. The RNA was eluted off the beads by treating them twice with 100 µl pre-heated (80°C) 100 mM β-mercaptoethanol (Merck, Cat#M6250), which breaks the disulfide bond between Biotin and 4sU. Subsequently, the RNA was converted into a deep sequencing library by following the manual instructions (Rev. C, 8/2014) of the ScriptSeq v2 RNA-Seq Library Preparation Kit (Illumina, Cat#SSV21106) starting with 4.1.A. (Anneal the cDNA Synthesis Primer) and 4.1.B. (Synthesize cDNA), RNA and ending with part 3.C (Synthesize 3’-Tagged DNA) to 3.G. (Assess Library Quantity and Quality). cDNA was purified using 1.8× SPRI beads. Input and 4sU-enriched cDNA were PCR-amplified with 11 and 15 cycles, respectively. The 4sU RNA-Seq library was purified with 1× SPRI beads.

#### Post-Sequencing Analysis of 4sU Tagging

Paired-end reads were aligned to the *X. tropicalis* transcriptome assembly v7.1 running Bowtie2 (Langmead and Salzberg, 2012) with the following constraints: -k 200 (maximal allowed number of alignments per fragment) -X 800 (maximum fragment length in bp) --rdg 6,5 (penalty for read gaps of length N, 6+N*5) --rfg 6,5 (penalty for reference gaps of length N, 6+N*5) --score-main L,-.6,-.4 (minimal alignment score as a linear function of the read length x, f(x) = −0.6 - 0.4*x) - -no-discordant (no paired-end read alignments breaching maximum fragment length X) --no-mixed (only concordant alignment of paired-end reads). Only read pairs that uniquely align to one gene were counted. Raw read counts were normalized with DESeq2 v1.22.1 (Love et al., 2014) and then scaled to the input.

#### Poly(A) RNA-Seq Profiling

10-15 embryos were collected per stage and condition. Total RNA was extracted as outlined above. Libraries were made from ∼1 µg total RNA by following the low-sample protocol of the TruSeq RNA Library Prep Kit v2 (Illumina, Cat#RS-122-2001) with a few modifications. First, 1 µl cDNA purified after second strand synthesis was quantified on a Qubit fluorometer using high-sensitivity reagents for detecting double-stranded DNA (10 pg/µl to 100 ng/µl). By this stage, the yield was ∼10 ng. Second, the number of PCR cycles was reduced to eight to avoid products of over-amplification such as chimera fragments.

#### Poly(A) RNA-Seq Read Alignment

Paired-end reads were aligned to the *X. tropicalis* genome assembly v7.1 using STAR v2.5.3a (Dobin et al., 2013) with default settings. The alignment was guided by a revised version of the gene models v7.2 (Collart et al., 2014) to improve mapping accuracy across splice junctions. The alignments were sorted by read name using the sort function of Samtools v1.3.1 (Li et al., 2009). Exon and intron counts (-t ‘exon;intron’) were extracted from unstranded (-s 0) alignment files using VERSE v0.1.5 (Zhu et al., 2016) in featureCounts (default) mode (-z 0). Intron coordinates were adjusted to exclude any overlap with exon annotation. For visualization, genomic BAM files of biological replicates were merged using Samtools and converted to the bigWig format. These genome tracks were normalized to the wigsum of 1 billion excluding any reads with mapping quality <10 using the python script bam2wig.py from RSeQC v2.6.4 (Wang et al., 2012).

#### Differential Gene Expression Analysis

Differential expression analysis was performed with both raw exon and intron counts excluding those belonging to ribosomal and mitochondrial RNA using the Bioconductor/R package DESeq2 v1.22.1 (Love et al., 2014). In an effort to find genes with consistent fold changes over time, p-values were generated according to a likelihood ratio test reflecting the probability of rejecting the reduced (∼ developmental stage) over the full (∼ developmental stage + condition) model. Resulting p-values were adjusted to obtain false discovery rates (FDR) according to the Benjamini-Hochburg procedure with thresholds on Cook’s distances and independent filtering being switched off. Equally, combinatorial LOF profiling and regional expression datasets (Blitz et al., 2017) without time series were subjected to likelihood ratio tests with reduced (∼ 1) and full (∼ condition) models for statistical analysis. Fold changes of intronic and exonic transcript levels were calculated for each developmental stage and condition using the mean of DESeq2-normalized read counts from biological replicates. Both intronic and exonic datasets were filtered for ≥10 DESeq2-normalized read counts that were detected at least at one developmental stage in all uninjected or DMSO-treated samples. Gene-specific fold changes were removed at developmental stages that yielded <10 normalized read counts in corresponding control samples. Next, the means of intronic and exonic fold changes were calculated across developmental stages. The whole dataset was confined to 3,318 genes for which at least 50% reductions (FDR ≤10%) in exonic (default) or intronic counts could be detected in α-amanitin-injected embryos. Regional expression was based on exonic read counts by default unless the intronic fold changes were significantly (FDR ≤10%) larger than the exonic fold changes (Table S3). For the hierarchical clustering of relative gene expression (Figure 2C), increased transcript levels were masked and only data points from signal LOFs, mPou5f3/Sox3 LOF and 4x zT LOF embryos were used. Euclidean distance-derived clusters were linked according to Ward’s criterion and sorted using the optimal leaf ordering (OLO) algorithm. The synergy factor (SF) between signals x and y (Figures 3 and S3) were calculated as follows: SF_xy_ = Δ_xy_ / (Δ_x_ + Δ_y_). Δ is the relative loss of gene expression caused by signal depletion. For these calculations, any gene upregulations were neutralised (i.e. set to 1).

#### Analysis of Enriched Gene Ontology (GO) Terms

Over-represented GO terms were found by applying hypergeometric tests of the Bioconductor/R package GOstats v2.42.0 (Falcon and Gentleman, 2007) on gene lists. The process was also supported by the Bioconductor/R packages GSEABase v1.44.0 (Morgan et al., 2017) and GO.db v3.4.1 (Carlson et al., 2007). The gene universe was associated with GO terms by means of BLAST2GO (Conesa et al., 2005) as previously outlined (Collart et al., 2014; Gentsch et al., 2015).

#### Generation of Hybridization Probes

Plasmids *X. laevis eomes* pCRII-TOPO (Gentsch et al., 2013) and *X. laevis tbxt (Xbra, t)* pSP73 (Smith et al., 1991) were linearized by restriction digestion (*Bam*HI and *Bgl*II, respectively) and purified using the QIAquick PCR Purification Kit (Qiagen, Cat#28104). The hybridization probes were transcribed from ∼1 μg linearized plasmid using 1× digoxigenin-11-UTP (Roche, Cat#11277065910), 40 U RiboLock RNase inhibitor (Thermo Fisher Scientific, Cat#EO0381), 1× transcription buffer (Roche) and T7 RNA polymerase (Roche, Cat#10881767001) at 37°C for 2 h. The probe was treated with 2 U Turbo DNase (Thermo Fisher Scientific) to remove the DNA template and purified by LiCl precipitation. RNA was diluted to 10 ng/μl (10× stock) with hybridization buffer. The hybridization buffer (stored at −20°C) consists of 50% formamide (Fisher Scientific, Cat#10052370), 5× saline sodium citrate (SSC), 1× Denhardt’s solution (Thermo Fisher Scientific, Cat#750018), 10 mM EDTA, 1 mg/ml torula RNA (Merck, Cat#R6625), 100 μg/ml heparin (Merck, Cat#H4784), 0.1% (v/v) Tween-20 (Merck, Cat#P9416) and 0.1% (w/v) CHAPS (Merck, Cat#C3023).

#### Whole-Mount In Situ Hybridization (WMISH)

WMISH was conducted using digoxigenin-labeled RNA probes (Monsoro-Burq, 2007; Sive et al., 2000). Briefly, *X. tropicalis* embryos were fixed in MEMFA (100 mM MOPS pH 7.4, 2 mM EDTA, 1 mM MgSO_4_ and 3.7% formaldehyde) at room temperature for 1 h. The embryos were then washed once in 1x PBS and two to three times in ethanol. Fixed and dehydrated embryos were kept at −20°C for ≥24 h to ensure proper dehydration before starting hybridization. Dehydrated embryos were washed once more in ethanol before rehydrating them in two steps to PBT (1×PBS and 0.1% (v/v) Tween-20). Embryos were treated with 5 μg/ml proteinase K (Thermo Fisher Scientific) in PBT for 6-8 min, washed briefly in PBT, fixed again in MEMFA for 20 min and washed three times in PBT. Embryos were transferred into baskets, which were kept in an 8×8 microcentrifuge tube holder sitting inside a 10×10 slot plastic box filled with PBT. Baskets were built by replacing the round bottom of 2-ml microcentrifuge tubes with a Sefar Nitex mesh. This container system was used to readily process several batches of embryos at once. These baskets were maximally loaded with 40 to 50 *X. tropicalis* embryos. The microcentrifuge tube holder was used to transfer all baskets at once and to submerge embryos into subsequent buffers of the WMISH protocol. Next, the embryos were incubated in 500 μl hybridization buffer (see recipe above) for 2 h in a hybridization oven set to 60°C. After this pre-hybridization step, the embryos were transferred into 500 μl digoxigenin-labeled probe (1 ng/µl) preheated to 60°C and further incubated overnight at 60°C. The pre-hybridization buffer was kept at 60°C. The next day embryos were transferred back into the pre-hybridization buffer and incubated at 60°C for 10 min. Subsequently, they were washed three times in 2× SSC/0.1% Tween-20 at 60°C for 10 min, twice in 0.2x SSC/0.1% Tween-20 at 60°C for 20 min and twice in 1x maleic acid buffer (MAB) at room temperature for 5 min. Next, the embryos were treated with blocking solution (2% Blocking Reagent [Merck, Cat#11096176001] in 1× MAB) at room temperature for 30 min, and incubated in antibody solution (10% lamb serum [Thermo Fisher Scientific, Cat#16070096], 2% Blocking Reagent [Merck], 1× MAB and 1:2,000 Fab fragments from polyclonal anti-digoxigenin antibodies conjugated to alkaline phosphatase [Roche, Cat#11093274910; RRID:AB_514497]) at room temperature for 4 h. The embryos were then washed four times in 1× MAB for 10 min before leaving them in 1× MAB overnight at 4°C.

On the final day of the WMISH protocol, the embryos were washed another three times in 1x MAB for 20 min and equilibrated to working conditions of alkaline phosphatase (AP) for a total of 10 min by submerging embryos twice into AP buffer (50 mM MgCl_2_, 100 mM NaCl, 100 mM Tris-HCl pH 9.5 and 1% (v/v) Tween-20). At this stage, the embryos were transferred to 5-ml glass vials for monitoring the progression of the AP-catalyzed colorimetric reaction. Any residual AP buffer was discarded before adding 700 μl staining solution (AP buffer, 340 μg/ml nitro-blue tetrazolium chloride [Roche, Cat#11383213001] and 175 μg/ml 5-bromo-4-chloro-3’-indolyphosphate [Roche, Cat#11383221001]). The colorimetric reaction was developed at room temperature in the dark. Once the staining was clear and intense enough, the color reaction was stopped by two washes in 1x MAB. To stabilize and preserve morphological features, the embryos were fixed with Bouin’s fixative without picric acid (9% formaldehyde and 5% glacial acetic acid [Fisher Scientific, Cat#10171460]) at room temperature for 30 min. Next, the embryos were washed twice in 70% ethanol/PBT to remove the fixative and residual chromogens. After rehydration to PBT in two steps, the embryos were treated with weak Curis solution (1% (v/v) hydrogen peroxide [Merck, Cat#1072090500], 0.5x SSC and 5% formamide) at 4°C in the dark overnight. Finally, the embryos were washed twice in PBS before imaging them in PBS on a thick agarose dish by light microscopy.

#### Processing of External Datasets

High-time (30-min intervals) resolution of total and poly(A) RNA-Seq (GSE65785) was processed as reported in the original publication (Owens et al., 2016). In addition, intron read counts were corrected by spike RNA-derived normalization factors. For visualization normalized exon and intron counts were scaled to the maximal count detected across the time course and fitted using cubic smoothing splines from 0 to 23.5 hpf: smooth.spline(1:48, ×, spar=0.6). Other RNA-Seq (GSE81458) and ChIP-Seq (GSE67974, GSE30146, GSE53654 and GSE72657) were processed as described in detail above except for H3K4me3 and H3K36me3 whose enriched regions were detected as follows: findPeaks -style histone - fragLength 175 -inputFragLength 175 -fdr 0.001 -gsize 1.435e9 -F 2 -C 1 -region -size 350 - minDist 500. Thus, we detected significant regions of histone modifications (-style histone) of at least the lengths of two DNA fragments (-size 350) and being separated by at least 500 bp from each other.

#### Generation of Plots and Heatmaps

Genomic snapshots were generated with the IGV genome browser v2.4-rc6 (Robinson et al., 2011). All plots and heatmaps were generated using R v3.5.1 (http://cran.r-project.org/). The following add-on R and Bioconductor packages were used for sorting and graphical visualization of data: alluvial v0.1-2 (Michal Bojanowsk), beeswarm v0.2.3 (Aron Eklund), circlize v0.4.5 (Gu et al., 2014), complexHeatmap v1.20.0 (Gu et al., 2016a), dplyr v0.7.8, ggplot2 v3.1.0 (Wickham, 2016), gplots v3.0.1 (Gregory Warnes and colleagues), GenomicFeatures v1.34.1 (Lawrence et al., 2013), GenomicRanges v1.38.0 (Lawrence et al., 2013), HilbertCurve v1.12.0 (Gu et al., 2016b), limma v3.38.2 (Ritchie et al., 2015), rtracklayer 1.42.1 (Lawrence et al., 2009) and seriation v1.2-3 (Hahsler et al., 2008).

### QUANTIFICATION AND STATISTICAL ANALYSIS

No statistical method was used for determining sample size; rather, we followed the literature to select the appropriate sample size. The experiments were not randomized. Due to the nature of experiments, the authors were not blinded to group allocation during data collection and analysis. Only viable embryos were included in the analysis. Frequencies of shown morphological phenotypes and WMISH patterns are included in every image. The significance of over-represented GO terms was based on hypergeometric tests. Significances of non-normally distributed data points (gene features) across ZGA were calculated using paired Wilcoxon rank-sum tests (alternative hypothesis ‘less’). The effect size (r_effect_) was estimated from the standard normal deviate of the Wilcoxon p-value (p) as previously described (Rosenthal, 1991), r_effect_=Z/sqrt(N), where Z=qnorm(1-p/2) is the standardized Z-score and N is the number of observations.

For RNA-Seq, biological triplicates were used to account for transcriptional variability between clutches. Each LOF experiment has its own control embryos collected in parallel from the same mothers: exp. #1 (α-amanitin), uninjected embryos; exp. #2 (BMP or Nodal LOF), DMSO-treated embryos; exp. #3 (Wnt LOF), uninjected embryos; exp. #4 (mPou5f3/Sox3 LOF), uninjected embryos; exp. #5 (mVegT LOF), uninjected embryos; exp. #6 (single and combinatorial LOFs of Wnt, Nodal and BMP), DMSO-treated embryos; exp. #7 (combinatorial LOF of 4 zygotic T-box TFs), control MO-injected embryos. The gene expressions of control MO-injected embryos of exp. #2 and #5 were normalized to their corresponding uninjected embryos. The mean of these normalizations and conservative FDR estimations (i.e., higher FDR of the two likelihood ratio tests) were used for the comparison with LOF conditions. RNA-Seq libraries from each experiment were generated simultaneously to mitigate any batch effects. The FDR was controlled for multiple comparisons according to the Benjamini-Hochberg procedure. The exact computational implementation of differential expression analysis is outlined on GitHub (see below).

### DATA AND SOFTWARE AVAILABILITY

Sequencing reads (FASTQ files) and raw RNA-Seq read counts reported in this paper are available in the GEO database (www.ncbi.nlm.nih.gov/geo) under the accession numbers GSE113186 and GSE122551. All analyses were performed in R v3.5.1 (Bioconductor v3.8), Perl v5.18.2 (https://www.perl.org) and Python v2.7.12 (http://www.python.org) as detailed above. The R code, genome annotation, intermediate datasets and graphs are available on GitHub at https://github.com/gegentsch/SpatioTemporalControlZGA. Original datasets are also available on Mendeley Data at http://dx.doi.org/10.17632/jn466b4n8v.1.

## REFERENCES

Agius, E., Oelgeschlager, M., Wessely, O., Kemp, C., and De Robertis, E.M. (2000). Endodermal Nodal-related signals and mesoderm induction in Xenopus. Development 127, 1173–1183.

Ali-Murthy, Z., Lott, S.E., Eisen, M.B., and Kornberg, T.B. (2013). An essential role for zygotic expression in the pre-cellular Drosophila embryo. PLoS Genet. 9, e1003428.

Anderson, G.A., Gelens, L., Baker, J.C., and Ferrell, J.E. (2017). Desynchronizing Embryonic Cell Division Waves Reveals the Robustness of Xenopus laevis Development. Cell Rep. 21, 37–46.

Arnold, S.J., and Robertson, E.J. (2009). Making a commitment: cell lineage allocation and axis patterning in the early mouse embryo. Nat. Rev. Mol. Cell Biol. 10, 91–103.

Boija, A., Klein, I.A., Sabari, B.R., Dall’Agnese, A., Coffey, E.L., Zamudio, A.V., Li, C.H., Shrinivas, K., Manteiga, J.C., Hannett, N.M., et al. (2018). Transcription Factors Activate Genes through the Phase-Separation Capacity of Their Activation Domains. Cell 175, 1842–1855.e16.

Blitz, I.L., Paraiso, K.D., Patrushev, I., Chiu, W.T.Y., Cho, K.W.Y., and Gilchrist, M.J. (2017). A catalog of Xenopus tropicalis transcription factors and their regional expression in the early gastrula stage embryo. Dev. Biol. 426, 409–417.

Blythe, S.A., and Wieschaus, E.F. (2015). Zygotic genome activation triggers the DNA replication checkpoint at the midblastula transition. Cell 160, 1169–1181.

Bolton, V.N., Oades, P.J., and Johnson, M.H. (1984). The relationship between cleavage, DNA replication, and gene expression in the mouse 2-cell embryo. J. Embryol. Exp. Morph. 79, 139–163.

Braude, P., Bolton, V., and Moore, S. (1988). Human gene expression first occurs between the four- and eight-cell stages of preimplantation development. Nature 332, 459–461.

Chan, S.H., Tang, Y., Miao, L., Darwich-Codore, H., Vejnar, C.E., Beaudoin, J.-D., Musaev, D., Fernandez, J.P., Moreno-Mateos, M.A., and Giraldez, A.J. (2018). Brd4 and P300 regulate zygotic genome activation through histone acetylation. bioRxiv 369231. DOI: https://doi.org/10.1101/369231

Chen, K., Hu, Z., Xia, Z., Zhao, D., Li, W., and Tyler, J.K. (2015). The Overlooked Fact: Fundamental Need for Spike-In Control for Virtually All Genome-Wide Analyses. Mol. Cell. Biol. 36, 662–667.

Chen, K., Johnston, J., Shao, W., Meier, S., Staber, C., and Zeitlinger, J. (2013). A global change in RNA polymerase II pausing during the Drosophila midblastula transition. Elife 2, e00861.

Chiu, W.T., Le, R.C., Blitz, I.L., Fish, M.B., Li, Y., Biesinger, J., Xie, X., and Cho, K.W.Y. (2014). Genome-wide view of TGFβ/Foxh1 regulation of the early mesendoderm program. Development 141, 1–114.

Cho, W.-K., Spille, J.-H., Hecht, M., Lee, C., Li, C., Grube, V., and Cisse, I.I. (2018). Mediator and RNA polymerase II clusters associate in transcription-dependent condensates. Science 361, 412–415.

Chong, S., Dugast-Darzacq, C., Liu, Z., Dong, P., Dailey, G.M., Cattoglio, C., Heckert, A., Banala, S., Lavis, L., Darzacq, X., et al. (2018). Imaging dynamic and selective low-complexity domain interactions that control gene transcription. Science 361, eaar2555.

Christian, J.L., and Moon, R.T. (1993). Interactions between Xwnt-8 and Spemann organizer signaling pathways generate dorsoventral pattern in the embryonic mesoderm of Xenopus. Genes Dev. 7, 13–28.

Collart, C., Owens, N.D.L., Bhaw-Rosun, L., Cooper, B., De Domenico, E., Patrushev, I., Sesay, A.K., Smith, J.N., Smith, J.C., and Gilchrist, M.J. (2014). High-resolution analysis of gene activity during the Xenopus mid-blastula transition. Development 141, 1927–1939.

Cuny, G.D., Yu, P.B., Laha, J.K., Xing, X., Liu, J.-F., Lai, C.S., Deng, D.Y., Sachidanandan, C., Bloch, K.D., and Peterson, R.T. (2008). Structure-activity relationship study of bone morphogenetic protein (BMP) signaling inhibitors. Bioorg. Med. Chem. Lett. 18, 4388–4392.

De Iaco, A., Coudray, A., Duc, J., and Trono, D. (2019). DPPA2 and DPPA4 are necessary to establish a 2C-like state in mouse embryonic stem cells. EMBO Reports e47382.

Du, Z., Zheng, H., Huang, B., Ma, R., Wu, J., Zhang, X., He, J., Xiang, Y., Wang, Q., Li, Y., et al. (2017). Allelic reprogramming of 3D chromatin architecture during early mammalian development. Nature 547, 232–235.

Eckersley-Maslin, M., Alda-Catalinas, C., Blotenburg, M., Kreibich, E., Krueger, C., and Reik, W. (2019). Dppa2 and Dppa4 directly regulate the Dux-driven zygotic transcriptional program. Genes Dev. 33, 194–208.

Faure, S., Lee, M.A., Keller, T., ten Dijke, P., and Whitman, M. (2000). Endogenous patterns of TGFbeta superfamily signaling during early Xenopus development. Development 127, 2917–2931.

Flyamer, I.M., Gassler, J., Imakaev, M., Brandão, H.B., Ulianov, S.V., Abdennur, N., Razin, S.V., Mirny, L.A., and Tachibana-Konwalski, K. (2017). Single-nucleus Hi-C reveals unique chromatin reorganization at oocyte-to-zygote transition. Nature 544, 110–114.

Fuentealba, L.C., Eivers, E., Ikeda, A., Hurtado, C., Kuroda, H., Pera, E.M., and De Robertis, E.M. (2007). Integrating patterning signals: Wnt/GSK3 regulates the duration of the BMP/Smad1 signal. Cell 131, 980–993.

Fukaya, T., Lim, B., and Levine, M. (2017). Rapid Rates of Pol II Elongation in the Drosophila Embryo. Curr. Biol. 27, 1387–1391.

Gentsch, G.E., Spruce, T., Monteiro, R.S., Owens, N.D.L., Martin, S.R., and Smith, J.C. (2018a). Innate Immune Response and Off-Target Mis-splicing Are Common Morpholino-Induced Side Effects in Xenopus. Dev. Cell 44, 597–610.e10.

Gentsch, G.E., Spruce, T., Owens, N.D.L., and Smith, J.C. (2018b). The role of maternal pioneer factors in predefining first zygotic responses to inductive signals. bioRxiv 306803. DOI: https://doi.org/10.1101/306803

Giraldez, A.J., Cinalli, R.M., Glasner, M.E., Enright, A.J., Thomson, J.M., Baskerville, S., Hammond, S.M., Bartel, D.P., and Schier, A.F. (2005). MicroRNAs regulate brain morphogenesis in zebrafish. Science 308, 833–838.

Giraldez, A.J., Mishima, Y., Rihel, J., Grocock, R.J., Van Dongen, S., Inoue, K., Enright, A.J., and Schier, A.F. (2006). Zebrafish MiR-430 promotes deadenylation and clearance of maternal mRNAs. Science 312, 75–79.

Gu, Z., Eils, R., and Schlesner, M. (2016). HilbertCurve: an R/Bioconductor package for high-resolution visualization of genomic data. Bioinformatics 32, 2372–2374.

Halley-Stott, R.P., Jullien, J., Pasque, V., and Gurdon, J. (2014). Mitosis gives a brief window of opportunity for a change in gene transcription. PLoS Biol. 12, e1001914.

Hamatani, T., Carter, M.G., Sharov, A.A., and Ko, M.S.H. (2004). Dynamics of global gene expression changes during mouse preimplantation development. Dev. Cell 6, 117–131.

Harvey, S.A., Sealy, I., Kettleborough, R., Fenyes, F., White, R., Stemple, D., and Smith, J.C. (2013). Identification of the zebrafish maternal and paternal transcriptomes. Development 140, 2703–2710.

Harvey, S.A., Tümpel, S., Dubrulle, J., Schier, A.F., and Smith, J.C. (2010). no tail integrates two modes of mesoderm induction. Development 137, 1127–1135.

Heasman, J. (2006). Patterning the early Xenopus embryo. Development 133, 1205–1217.

Heasman, J., Kofron, M., and Wylie, C. (2000). βCatenin Signaling Activity Dissected in the Early Xenopus Embryo: A Novel Antisense Approach. Dev. Biol. 222, 124–134.

Heyn, P., Kircher, M., Dahl, A., Kelso, J., Tomancak, P., Kalinka, A.T., and Neugebauer, K.M. (2014). The earliest transcribed zygotic genes are short, newly evolved, and different across species. Cell Rep. 6, 285–292.

Ho, D.M., Chan, J., Bayliss, P., and Whitman, M. (2006). Inhibitor-resistant type I receptors reveal specific requirements for TGF-beta signaling in vivo. Dev. Biol. 295, 730–742.

Hontelez, S., van Kruijsbergen, I., Georgiou, G., van Heeringen, S.J., Bogdanović, O., Lister, R., and Veenstra, G.J.C. (2015). Embryonic transcription is controlled by maternally definedchromatin state. Nat. Commun. 6, 10148.

Hug, C.B., Grimaldi, A.G., Kruse, K., and Vaquerizas, J.M. (2017). Chromatin Architecture Emerges during Zygotic Genome Activation Independent of Transcription. Cell 169, 216–228.e219.

Inman, G.J., Nicolás, F.J., Callahan, J.F., Harling, J.D., Gaster, L.M., Reith, A.D., Laping, N.J., and Hill, C.S. (2002). SB-431542 is a potent and specific inhibitor of transforming growth factor-beta superfamily type I activin receptor-like kinase (ALK) receptors ALK4, ALK5, and ALK7. Mol. Pharmacol. 62, 65–74.

Jones, C.M., Kuehn, M.R., Hogan, B.L., Smith, J.C., and Wright, C.V. (1995). Nodal-related signals induce axial mesoderm and dorsalize mesoderm during gastrulation. Development 121, 3651–3662.

Joseph, S.R., Pálfy, M., Hilbert, L., Kumar, M., Karschau, J., Zaburdaev, V., Shevchenko, A., and Vastenhouw, N.L. (2017). Competition between histone and transcription factor binding regulates the onset of transcription in zebrafish embryos. Elife 6, e23326.

Jukam, D., Shariati, S.A.M., and Skotheim, J.M. (2017). Zygotic Genome Activation in Vertebrates. Dev. Cell 42, 316–332.

Kaaij, L.J.T., van der Weide, R.H., Ketting, R.F., and de Wit, E. (2018). Systemic Loss and Gain of Chromatin Architecture throughout Zebrafish Development. Cell Rep. 24, 1–10.e4.

Kane, D.A., and Kimmel, C.B. (1993). The zebrafish midblastula transition. Development 119, 447–456.

Ke, Y., Xu, Y., Chen, X., Feng, S., Liu, Z., Sun, Y., Yao, X., Li, F., Zhu, W., Gao, L., et al. (2017). 3D Chromatin Structures of Mature Gametes and Structural Reprogramming during Mammalian Embryogenesis. Cell 170, 367–381.e20.

Kimelman, D. (2006). Mesoderm induction: from caps to chips. Nat. Rev. Genet. 7, 360–372.

Langley, A.R., Smith, J.C., Stemple, D.L., and Harvey, S.A. (2014). New insights into the maternal to zygotic transition. Development 141, 3834–3841.

Larabell, C.A., Torres, M., Rowning, B.A., Yost, C., Miller, J.R., Wu, M., Kimelman, D., and Moon, R.T. (1997). Establishment of the dorso-ventral axis in Xenopus embryos is presaged by early asymmetries in beta-catenin that are modulated by the Wnt signaling pathway. J. Cell Biol. 136, 1123–1136.

Lemaire, P., Garrett, N., and Gurdon, J.B. (1995). Expression cloning of Siamois, a Xenopus homeobox gene expressed in dorsal-vegetal cells of blastulae and able to induce a complete secondary axis. Cell 81, 85–94.

Lee, M.T., Bonneau, A.R., Takacs, C.M., Bazzini, A.A., Divito, K.R., Fleming, E.S., and Giraldez, A.J. (2013). Nanog, Pou5f1 and SoxB1 activate zygotic gene expression during the maternal-to-zygotic transition. Nature 503, 360–364.

Leichsenring, M., Maes, J., Mössner, R., Driever, W., and Onichtchouk, D. (2013). Pou5f1 transcription factor controls zygotic gene activation in vertebrates. Science 341, 1005–1009.

Liang, H.-L., Nien, C.-Y., Liu, H.-Y., Metzstein, M.M., Kirov, N., and Rushlow, C. (2008). The zinc-finger protein Zelda is a key activator of the early zygotic genome in Drosophila. Nature 456, 400–403.

Liu, G., Wang, W., Hu, S., Wang, X., and Zhang, Y. (2018). Inherited DNA methylation primes the establishment of accessible chromatin during genome activation. Genome Res. 28, 998–1007.

Lott, S.E., Villalta, J.E., Schroth, G.P., Luo, S., Tonkin, L.A., and Eisen, M.B. (2011). Noncanonical compensation of zygotic X transcription in early Drosophila melanogaster development revealed through single-embryo RNA-seq. PLoS Biol. 9, e1000590.

Lu, F., Liu, Y., Inoue, A., Suzuki, T., Zhao, K., and Zhang, Y. (2016). Establishing Chromatin Regulatory Landscape during Mouse Preimplantation Development. Cell 165, 1375–1388.

Lu, X., Li, J.M., Elemento, O., Tavazoie, S., and Wieschaus, E.F. (2009). Coupling of zygotic transcription to mitotic control at the Drosophila mid-blastula transition. Development 136, 2101–2110.

Lund, E., Liu, M., Hartley, R.S., Sheets, M.D., and Dahlberg, J.E. (2009). Deadenylation of maternal mRNAs mediated by miR-427 in Xenopus laevis embryos. RNA 15, 2351–2363.

Mathavan, S., Lee, S.G.P., Mak, A., Miller, L.D., Murthy, K.R.K., Govindarajan, K.R., Tong, Y., Wu, Y.L., Lam, S.H., Yang, H., et al. (2005). Transcriptome analysis of zebrafish embryogenesis using microarrays. PLoS Genet. 1, 260–276.

Newport, J., and Kirschner, M.W. (1982a). A major developmental transition in early xenopus embryos: I. characterization and timing of cellular changes at the midblastula stage. Cell 30, 675–686.

Newport, J., and Kirschner, M.W. (1982b). A major developmental transition in early xenopus embryos: II. control of the onset of transcription. Cell 30, 687–696.

Nudelman, G., Frasca, A., Kent, B., Sadler, K.C., Sealfon, S.C., Walsh, M.J., and Zaslavsky, E. (2018). High resolution annotation of zebrafish transcriptome using long-read sequencing. Genome Res. 28, 1415–1425.

Owens, N.D.L., Blitz, I.L., Lane, M.A., Patrushev, I., Overton, J.D., Gilchrist, M.J., Cho, K.W.Y., and Khokha, M.K. (2016). Measuring Absolute RNA Copy Numbers at High Temporal Resolution Reveals Transcriptome Kinetics in Development. Cell Rep. 14, 632–647.

Reversade, B., Kuroda, H., Lee, H., Mays, A., and De Robertis, E.M. (2005). Depletion of Bmp2, Bmp4, Bmp7 and Spemann organizer signals induces massive brain formation in Xenopus embryos. Development 132, 3381–3392.

Sabari, B.R., Dall’Agnese, A., Boija, A., Klein, I.A., Coffey, E.L., Shrinivas, K., Abraham, B.J., Hannett, N.M., Zamudio, A.V., Manteiga, J.C., et al. (2018). Coactivator condensation at super-enhancers links phase separation and gene control. Science 361, eaar3958.

Schneider, S., Steinbeisser, H., Warga, R., and Hausen, P. (1996). Beta-catenin translocation into nuclei demarcates the dorsalizing centers in frog and fish embryos. Mech. Dev. 57, 191–198.

Schohl, A., and Fagotto, F. (2002). Beta-catenin, MAPK and Smad signaling during early Xenopus development. Development 129, 37–52.

Shermoen, A.W., and O’Farrell, P.H. (1991). Progression of the cell cycle through mitosis leads to abortion of nascent transcripts. Cell 67, 303–310.

Shrinivas, K., Sabari, B.R., Coffey, E.L., Klein, I.A., Boija, A., Zamudio, A.V., Schuijers, J., Hannett, N.M., Sharp, P.A., Young, R.A., et al. (2018). Enhancer features that drive formation of transcriptional condensates. bioRxiv 495606. DOI: https://doi.org/10.1101/495606

Skirkanich, J., Luxardi, G., Yang, J., Kodjabachian, L., and Klein, P.S. (2011). An essential role for transcription before the MBT in Xenopus laevis. Dev. Biol. 357, 478–491.

Stack, J.H., and Newport, J.W. (1997). Developmentally regulated activation of apoptosis early in Xenopus gastrulation results in cyclin A degradation during interphase of the cell cycle. Development 124, 3185–3195.

Takahashi, S., Onuma, Y., Yokota, C., Westmoreland, J.J., Asashima, M., and Wright, C.V.E. (2006). Nodal-related gene Xnr5 is amplified in the Xenopus genome. Genesis 44, 309–321.

Tan, M.H., Au, K.F., Yablonovitch, A.L., Wills, A.E., Chuang, J., Baker, J.C., Wong, W.H., and Li, J.B. (2013). RNA sequencing reveals a diverse and dynamic repertoire of the Xenopus tropicalis transcriptome over development. Genome Res. 23, 201–216.

Tani, S., Kusakabe, R., Naruse, K., Sakamoto, H., and Inoue, K. (2010). Genomic organization and embryonic expression of miR-430 in medaka (Oryzias latipes): insights into the post-transcriptional gene regulation in early development. Gene 449, 41–49.

Tao, Q., Yokota, C., Puck, H., Kofron, M., Birsoy, B., Yan, D., Asashima, M., Wylie, C.C., Lin, X., and Heasman, J. (2005). Maternal wnt11 activates the canonical wnt signaling pathway required for axis formation in Xenopus embryos. Cell 120, 857–871.

Vassena, R., Boué, S., González-Roca, E., Aran, B., Auer, H., Veiga, A., and Izpisúa Belmonte, J.C. (2011). Waves of early transcriptional activation and pluripotency program initiation during human preimplantation development. Development 138, 3699–3709.

Whyte, W.A., Orlando, D.A., Hnisz, D., Abraham, B.J., Lin, C.Y., Kagey, M.H., Rahl, P.B., Lee, T.I., and Young, R.A. (2013). Master transcription factors and mediator establish super-enhancers at key cell identity genes. Cell 153, 307–319.

Wu, J., Huang, B., Chen, H., Yin, Q., Liu, Y., Xiang, Y., Zhang, B., Liu, B., Wang, Q., Xia, W., et al. (2016). The landscape of accessible chromatin in mammalian preimplantation embryos. Nature 534, 652–657.

Yanai, I., Peshkin, L., Jorgensen, P., and Kirschner, M.W. (2011). Mapping gene expression in two Xenopus species: evolutionary constraints and developmental flexibility. Dev. Cell 20, 483–496.

Yang, J., Tan, C., Darken, R.S., Wilson, P.A., and Klein, P.S. (2002). Beta-catenin/Tcf-regulated transcription prior to the midblastula transition. Development 129, 5743–5752.

Young, J.J., Kjolby, R.A.S., Wu, G., Wong, D., Hsu, S.-W., and Harland, R.M. (2017). Noggin is required for first pharyngeal arch differentiation in the frog Xenopus tropicalis. Dev. Biol. 426, 245–254.

## SUPPLEMENTAL REFERENCES

Benson, G. (1999). Tandem repeats finder: a program to analyze DNA sequences. Nucleic Acids Res. 27, 573–580.

Carlson, M., Falcon, S., Pages, H., and Li, N. (2007). Bioconductor - GO.db.

Conesa, A., Götz, S., García-Gómez, J.M., Terol, J., Talón, M., and Robles, M. (2005). Blast2GO: a universal tool for annotation, visualization and analysis in functional genomics research. Bioinformatics 21, 3674–3676.

Dobin, A., Davis, C.A., Schlesinger, F., Drenkow, J., Zaleski, C., Jha, S., Batut, P., Chaisson, M., and Gingeras, T.R. (2013). STAR: ultrafast universal RNA-seq aligner. Bioinformatics 29, 15–21.

Falcon, S., and Gentleman, R. (2007). Using GOstats to test gene lists for GO term association. Bioinformatics 23, 257–258.

Fukuda, M., Takahashi, S., Haramoto, Y., Onuma, Y., Kim, Y.-J., Yeo, C.-Y., Ishiura, S., and Asashima, M. (2010). Zygotic VegT is required for Xenopus paraxial mesoderm formation and is regulated by Nodal signaling and Eomesodermin. Int. J. Dev. Biol. 54, 81–92.

Gay, L., Karfilis, K.V., Miller, M.R., Doe, C.Q., and Stankunas, K. (2014). Applying thiouracil tagging to mouse transcriptome analysis. Nat. Protoc. 9, 410–420.

Gentsch, G.E., and Smith, J.C. (2017). Efficient Preparation of High-Complexity ChIP-Seq Profiles from Early Xenopus Embryos. Methods Mol. Biol. 1507, 23–42.

Gentsch, G.E., Owens, N.D.L., Martin, S.R., Piccinelli, P., Faial, T., Trotter, M.W.B., Gilchrist, M.J., and Smith, J.C. (2013). In vivo T-box transcription factor profiling reveals joint regulation of embryonic neuromesodermal bipotency. Cell Rep. 4, 1185–1196.

Gentsch, G.E., Patrushev, I., and Smith, J.C. (2015). Genome-wide snapshot of chromatin regulators and states in Xenopus embryos by ChIP-Seq. J. Vis. Exp. 96, e52535.

Gu, Z. Eils, R., and Schlesner, M. (2016a). Complex heatmaps reveal patterns and correlations in multidimensional genomic data. Bioinformatics 32, 2847–2849.

Gu, Z., Eils, R., and Schlesner, M. (2016b). HilbertCurve: an R/Bioconductor package for high-resolution visualization of genomic data. Bioinformatics 32, 2372–2374.

Gu, Z., Gu, L., Eils, R., Schlesner, M., and Brors, B. (2014). circlize Implements and enhances circular visualization in R. Bioinformatics 30, 2811–2812.

Hahsler, M., Hornik, K., and Buchta, C. (2008). Getting Things in Order: An Introduction to the R Package seriation. J. Stat. Softw. 25, 1–34.

Heinz, S., Benner, C., Spann, N., Bertolino, E., Lin, Y.C., Laslo, P., Cheng, J.X., Murre, C., Singh, H., and Glass, C.K. (2010). Simple combinations of lineage-determining transcription factors prime cis-regulatory elements required for macrophage and B cell identities. Mol. Cell 38, 576–589.

Hontelez, S., van Kruijsbergen, I., Georgiou, G., van Heeringen, S.J., Bogdanović, O., Lister, R., and Veenstra, G.J.C. (2015). Embryonic transcription is controlled by maternally defined chromatin state. Nat. Commun. 6, 10148.

Khokha, M., Chung, C., Bustamante, E., Gaw, L., Trott, K., Yeh, J., Lim, N., Lin, J., Taverner, N., Amaya, E., et al. (2002). Techniques and probes for the study of Xenopus tropicalis development. Dev. Dyn. 225, 499–510.

Langmead, B., and Salzberg, S.L. (2012). Fast gapped-read alignment with Bowtie 2. Nat. Methods 9, 357–359.

Lawrence, M., Gentleman, R., and Carey, V. (2009). rtracklayer: an R package for interfacing with genome browsers. Bioinformatics 25, 1841–1842.

Lawrence, M., Huber, W., Pagè s, H., Aboyoun, P., Carlson, M., Gentleman, R., Morgan, M., and Carey, V. (2013). Software for Computing and Annotating Genomic Ranges. PLoS Comput. Biol. 9, e1003118.

Li, H., Handsaker, B., Wysoker, A., Fennell, T., Ruan, J., Homer, N., Marth, G., Abecasis, G., Durbin, R., 1000 Genome Project Data Processing Subgroup (2009). The Sequence Alignment/Map format and SAMtools. Bioinformatics 25, 2078–2079.

Love, M.I., Huber, W., and Anders, S. (2014). Moderated estimation of fold change and dispersion for RNA-seq data with DESeq2. Genome Biol. 15, 550.

Monsoro-Burq, A.H. (2007). A Rapid Protocol for Whole-Mount In Situ Hybridization on Xenopus Embryos. Cold Spring Harb. Protoc. 2007, pdb.prot4809.

Morgan, M., Falcon, S., and Gentleman, R. (2017). GSEABase: Gene set enrichment data structures and methods.

Nakamura, Y., de Paiva Alves, E., Veenstra, G.J.C., and Hoppler, S. (2016). Tissue- and stage-specific Wnt target gene expression is controlled subsequent to β-catenin recruitment to cis-regulatory modules. Development 143, 1914–1925.

Nieuwkoop, P., and Faber, J. (1994). Normal table of Xenopus laevis (Daudin): a systematical and chronological survey of the development from the fertilized egg till the end of metamorphosis. Garland.

Pertea, M., Kim, D., Pertea, G.M., Leek, J.T., and Salzberg, S.L. (2016). Transcript-level expression analysis of RNA-seq experiments with HISAT, StringTie and Ballgown. Nat. Protoc. 11, 1650–1667.

Quinlan, A.R., and Hall, I.M. (2010). BEDTools: a flexible suite of utilities for comparing genomic features. Bioinformatics 26, 841–842.

Rana, A., Collart, C., Gilchrist, M., and Smith, J. (2006). Defining synphenotype groups in Xenopus tropicalis by use of antisense morpholino oligonucleotides. PLoS Genet. 2, e193.

Ritchie, M.E., Phipson, B., Wu, D., Hu, Y., Law, C.W., Shi, W., and Smyth, G.K. (2015). limma powers differential expression analyses for RNA-sequencing and microarray studies. Nucleic Acids Res. 43, e47.

Robinson, J.T., Thorvaldsdóttir, H., Winckler, W., Guttman, M., Lander, E.S., Getz, G., and Mesirov, J.P. (2011). Integrative genomics viewer. Nat. Biotechnol. 29, 24–26.

Rosenthal, R. (1991). Meta-Analytic Procedures for Social Research. SAGE Publishing.

Sive, H., Grainger, R., and Harland, R. (2000). Early development of Xenopus laevis: A laboratory manual. Cold Spring Harbor Laboratory Press.

Smit, A., Hubley, R., and Green, P. RepeatMasker Open-4.0. 2013-2015.

Smith, J., Price, B., Green, J., Weigel, D., and Herrmann, B. (1991). Expression of a Xenopus homolog of Brachyury (T) is an immediate-early response to mesoderm induction. Cell 67, 79–87.

Wang, L., Wang, S., and Li, W. (2012). RSeQC: quality control of RNA-seq experiments. Bioinformatics 28, 2184–2185.

Wickham, H. (2016). ggplot2: Elegant Graphics for Data Analysis. Springer.

Yoon, S.-J., Wills, A.E., Chuong, E., Gupta, R., and Baker, J.C. (2011). HEB and E2A function as SMAD/FOXH1 cofactors. Genes Dev. 25, 1654–1661.

Zhu, Q., Fisher, S.A., Shallcross, J., and Kim, J. (2016). VERSE: a versatile and efficient RNA-Seq read counting tool. bioRxiv 053306. DOI: https://doi.org/10.1101/053306

